# Bioenergetic dysfunction and inflammation in hiPSC-derived astrocytes from m.14484T>C Leber’s Hereditary Optic Neuropathy

**DOI:** 10.64898/2026.07.23.740295

**Authors:** Wyn Firth, Lubica Dudakova, Robert Dobrovolny, Tomas Honzik, Petra Liskova, Julie Albon, Marcela Votruba

## Abstract

Leber’s Hereditary Optic Neuropathy (LHON) is a maternally inherited mitochondrial disorder characterised by painless, progressive, and sequential visual failure. Most cases of LHON are driven by mitochondrial DNA mutations which cause dysfunction of respiratory Complex I, triggering retinal ganglion cell loss. Retinal ganglion cell degeneration in LHON is thought to be linked to reduced production of metabolic intermediates and adenosine triphosphate, and enhanced reactive oxygen species production. Thus, decades of research have focussed on LHON as a disease of the retinal ganglion cells, which has considerably improved our understanding of the pathology but has yielded few therapeutic interventions. In addition, some LHON-associated phenomena remain unclear. In particular, we still do not fully understand the mechanisms underlying the recorded phenomenon of spontaneous visual recovery, in which patients experience measurable increases in visual acuity following onset of LHON vision loss. Understanding this phenomenon may be critical for developing new therapeutic approaches for LHON. Moreover, the contribution of non-neuronal cell populations to LHON pathology remains poorly understood despite a growing appreciation for the roles played by these cells in other neurodegenerative conditions. Astrocytes are a highly heterogeneous group of glial cells, found throughout the central nervous system including the retina and optic nerve, and are well-known for their role as key homeostatic mediators. In recent years, our appreciation for the role played by astrocytes in neurodegenerative diseases has expanded considerably, and we are now aware that astrocytes undergo significant loss of their homeostatic functions in neurodegenerative disease, acting as key mediators of neuronal loss. Importantly, the contributions astrocytes make toward mediating LHON pathology and visual recovery remain unclear, and provide promising ground for potentially novel therapeutic angles and enhanced understanding of this complex pathology. Here, we leverage human iPSC-derived astrocytes from patients with the LHON m.14484T>C genotype, to explore the role astrocytes play in LHON pathology, stratifying cells by their visual recovery status. We report that astrocytes undergo significant morphological and bioenergetic compromise in LHON, and that differences between ‘recovery’ and ‘non-recovery’ astrocytes may explain individual capacity for visual recovery, potentially opening novel therapeutic approaches.

## Introduction

Leber’s Hereditary Optic Neuropathy (LHON [OMIM #535000])^1^ is a maternally inherited neurodegenerative condition affecting ∼1/30,000-1/50,000 people worldwide^2,3^. In LHON, patients aged 2-87 years^4–6^ of both sexes experience painless, (sub)acute, bilateral vision loss resulting in central scotoma. LHON is classically considered to be driven by mutations to the mitochondrial genome; estimates suggest that >90% of LHON cases are driven by one of three point mutations^4,7,8^. In order of frequency these are m.11778G>A, m.14484T>C, and m.3460G>A^4,7,8^. Despite affecting different loci, these mutations have the common effect of driving respiratory Complex I dysfunction^9^. Common consensus suggests that, in LHON, Complex I dysfunction-associated reductions in cellular adenosine triphosphate (ATP) production and enhanced reactive oxygen species (ROS) production drive the death of retinal ganglion cells (RGCs), the output neurons of the retina. Crucially, despite many decades of research, therapeutic options for LHON remain limited. It has thus become clear that the contributions of other retinal cell populations to LHON pathology may further explain the underlying pathophysiological mechanisms of RGC loss and could open up avenues for the development of clinically useful therapeutic agents. An unusual feature of LHON among neurodegenerative diseases is the recorded capacity for some degree of spontaneous improvement in central vision^5,6,8^. The mechanistic basis for this unusual phenomenon remains unclear; however, it is known to occur most frequently in the m.14484T>C genotype^5,6,10^. This observation is highly intriguing as it suggests that there is potential for visual recovery, and if the mechanisms and drivers can be elucidated, potentially a new approach to LHON therapeutics.

Astrocytes are a highly heterogeneous class of glial cells found throughout the central nervous system (CNS), including the central retina^11^ and optic nerve^12–16^, which play multiple indispensable roles that have the net effect of maintaining CNS homeostasis in normal physiology. Changes to astrocyte physiology are now known to form a key part of various neurodegenerative conditions^16–23^, but the potential role of astrocytes in LHON pathology remains unclear. Crucially, astrocyte metabolism forms a nexus for various homeostatic functions including ion homeostasis, inflammatory responses, and providing trophic support to neurons. Bioenergetically, astrocytes are commonly regarded as showing greater reliance on aerobic glycolysis than other CNS cell types such as microglia and neurons, with a relatively lower level of oxidative phosphorylation (OXPHOS) than their counterparts^24^. Despite this, Complex I dysfunction is known to modulate the astrocyte phenotype and promote an inflammatory response, as seen in a model of Leigh syndrome^25^. This area of research is of growing importance for the neurodegenerative research community and reflects an increased interest in glial perturbations in other retinal neurodegenerative conditions such as glaucoma^16–18^. Indeed, changes to astrocyte metabolism are now known to influence the inflammatory responses of these cells, and shifts to astrocyte metabolism are associated with other conditions^19,26^. Critically, the role of astrocytes in LHON pathology, as well as the impact of LHON-specific mutations on astrocyte morphology, bioenergetics, and function, remain poorly understood despite being touted as a potentially fruitful avenue for novel LHON therapeutics^8^.

To explore LHON-associated changes to astrocyte form and function, here we leveraged LHON patient-derived induced pluripotent stem cells (iPSCs) from two unrelated donors ± spontaneous visual recovery (R: recovery, NR: non-recovery) carrying the m.14484T>C mutation, and an unrelated healthy control. These iPSCs were used to generate LHON-affected astrocytes stratified by visual recovery status to test the hypothesis that differential changes to astrocytes in LHON recovery states may contribute to capacity for spontaneous visual recovery.

## Methods

### Ethics statement

The study followed the tenets of the Declaration of Helsinki and was approved by the Ethics Committee of the General University Hospital (protocol code 25/18 Grant AZV VES 2019 VFN), under full informed consent allowing for secondary academic research and international transfer.

### General cell culture

All cultureware were coated with Geltrex (Fisher, cat. no. 15180617) prior to use. Geltrex was aspirated immediately prior to seeding and hydration maintained by the addition of media, if required.

### iPSC generation and culture

Induced pluripotent stem cells (iPSCs) were generated as described previously^27^ from two, unrelated, LHON-affected donors ± spontaneous visual recovery within 5 years from vision loss (hereinafter referred to as recovery [R, LHON-R] and non-recovery [NR, LHON-NR]) carrying the m.14484T>C mutation, and an unrelated healthy control. Briefly, peripheral blood mononuclear cells were isolated from LHON donors, and reprogrammed using the CytoTune-iPS 2.0 Sendai Reprogramming kit with a multiplicity of infection of 5:5:3 hKOS:hc-Myc:hKlf4. iPSCs were adapted to feeder-free culture and maintained as colonies in Geltrex-coated 6-well plates (Sarstedt, cat. no. 83.3920) in mTeSR Plus media (Stem Cell Technologies [SCT], cat. no. 100-1130). iPSCs underwent aggregate passage once weekly. Briefly, media was removed and cells were incubated in cell dissociation buffer (Fisher, cat. no. 13150016) for 2-3 minutes at 37°C. Colonies were monitored for signs of detachment. Cell dissociation buffer was aspirated and detachment facilitated by gentle rinsing with mTeSR Plus media supplemented with 10µM Y-27632 (SCT, cat. no. 72304; 0.01M PBS vehicle). Y-27632 was removed after 6-24h of culture to avoid adverse effects. Media was changed every 1-3 days, depending on confluency, and cells monitored for spontaneous differentiation. Spontaneously differentiated cells were identified by morphology and removed by scraping and aspiration. All cell lines were confirmed mycoplasma-free via the Lonza MycoAlert Plus Kit (Lonza, cat. no. LT37-701) prior to differentiations.

### Astrocyte differentiation

Astrocyte differentiations were performed using the Astrocyte Differentiation Kit (SCT, cat. no. 100-0013) according to the manufacturer’s specifications with minor modifications. All stages requiring adherent cells were performed using Geltrex-coated cultureware. Briefly, iPSCs were dissociated with accutase (SCT, cat. no. 07922; or Merck, cat. no. A6964) and seeded at 5×10^3^ cells per well in AggreWell 800 plates to form embryoid bodies according to the manufacturer’s instructions (Supplementary Figure 1A). Embryoid bodies were differentiated to a neural progenitor cell (NPC) lineage using the STEMdiff^TM^ Neural Induction Kit (SCT cat. no. 08582) (Supplementary Figure 1A). Following isolation, NPCs were differentiated into astrocyte progenitor cells using the STEMdiff^TM^ Astrocyte Differentiation Kit (SCT, cat. no. 100-0013) (Supplementary Figure 1A). Astrocyte progenitor cells were matured to terminally differentiated astrocytes using the STEMdiff^TM^ Astrocyte Serum-Free Maturation Kit (SCT, cat. no. 100-1666) (Supplementary Figure 1A). Terminally differentiated mature astrocytes were passaged once weekly, at a density of 1×10^5^ cells per well (LHON-affected cells) or 4×10^5^-1.44×10^6^ cells per well (unrelated healthy control cells), according to the manufacturer’s specifications with a minor modification – cells were incubated with 10µM Y-27632 for 2-6h post-seed to encourage viability. Cells were fed Astrocyte Serum-Free Maturation Media (ASFMM) every 2-3 days. Mature astrocytes were used for experiments from 60-90 days-post-differentiation (dpd; i.e., 60-90 days since embryoid body formation). Excess terminally differentiated astrocytes were maintained in ASFMM for a maximum of 10 passages (astrocytes were passaged as required, generally once every 1-2 weeks), or cryopreserved in CryoStor CS-10 (CST, cat. no. 100-1061). Cells were allowed a minimum of 1 week in ASFMM to recover from cryopreservation prior to use in assays. Astrocyte fate specification was determined via immunocytochemistry for the canonical *in vitro* astrocyte markers glial fibrillary acidic protein (GFAP) and S100 calcium binding protein B (S100B) (Supplementary Figure 1B,C); cultures were 100% positive for GFAP or S100B or both markers. All data presented herein are derived from mature astrocytes, 60-90 dpd.

### Seeding

Media was aspirated, cells washed once with DMEM/F12 (Fisher, cat. no. 11574546) and dissociated with accutase at 37°C 5% CO_2_ for 5 minutes. Accutase was neutralised by addition of an equal volume of DMEM/F12. Cells were gently triturated and vessels rinsed with DMEM/F12. Cells were centrifuged at 400*xg* for 5 minutes at room temperature and pressure (RTP) and resuspended in 1mL ASFMM. Cells were counted using a Countess III cell counter (Fisher) and seeded at the appropriate densities in seeding media (Table 1). After 16-24h, media was replaced with assay media for 2h prior to cell harvesting or use in experiments.

**Table 1:**
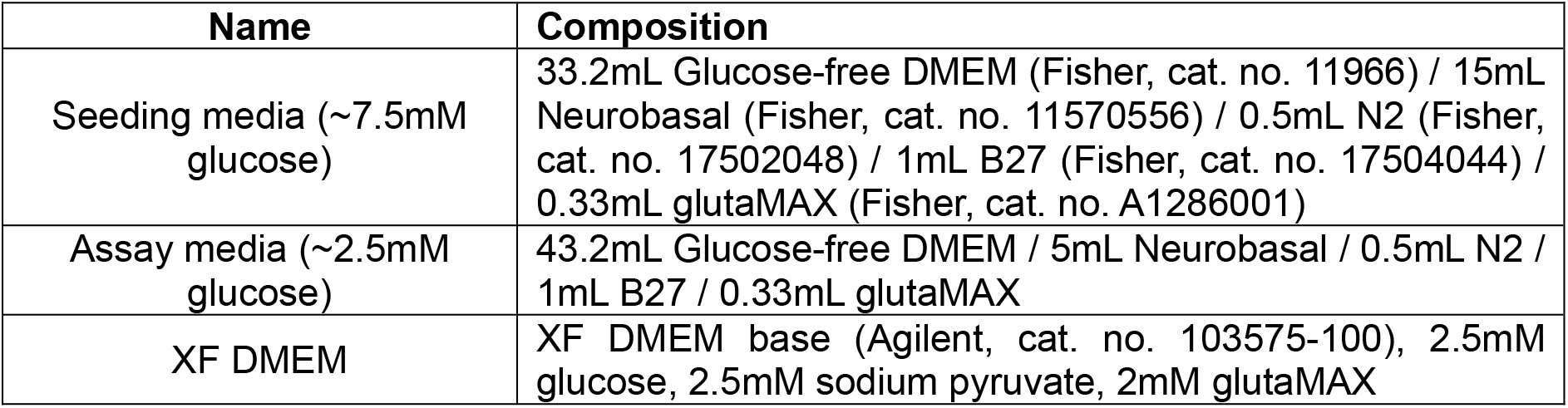
Media recipes.

| Name | Composition |
| --- | --- |
| Seeding media (~7.5mM glucose) | 33.2mL Glucose-free DMEM (Fisher, cat. no. 11966) / 15mL Neurobasal (Fisher, cat. no. 11570556) / 0.5mL N2 (Fisher, cat. no. 17502048) / 1mL B27 (Fisher, cat. no. 17504044) / 0.33mL glutaMAX (Fisher, cat. no. A1286001) |
| Assay media (~2.5mM glucose) | 43.2mL Glucose-free DMEM / 5mL Neurobasal / 0.5mL N2 / 1mL B27 / 0.33mL glutaMAX |
| XF DMEM | XF DMEM base (Agilent, cat. no. 103575-100), 2.5mM glucose, 2.5mM sodium pyruvate, 2mM glutaMAX |

### Extracellular metabolic flux analyses

Extracellular metabolic flux was measured using a Seahorse XFe96 bioanalyser (Agilent). Cells were plated 16-24h prior to the assay at a density of 2.0×10^4^ cells per well. Prior to all assays, assay media was removed and cells washed once with XF DMEM (Table 1). This media was immediately aspirated and replaced with fresh XF DMEM of the same composition. Cells were then placed into a humidified, 37°C, non-CO_2_ incubator to degas for 1 hour before immediately proceeding to assays. All assays were run at 37°C and within a consistent time window (assay commencement 1130-1200h) to control for circadian rhythmicity. A 3:0:3 mix:wait:measure ratio was used for all assays and 4 baseline reads were taken prior to any injections. Following all assays, XF DMEM was aspirated and cells lysed in 50mM NaOH. Plates were stored at −20°C for a maximum of 5 days. Protein content per well was estimated via a Bradford assay as described previously^28,29^ and extracellular metabolic flux data were normalised to protein content.

### Mitochondrial stress test

Mitochondrial function was gauged via a mitochondrial stress test (Agilent, cat. no. 103015-100). Compounds were reconstituted in XF DMEM (Table 1) and loaded into ports. Oligomycin (Oligo), carbonyl cyanide-p-trifluoromethoxyphenylhydrazone (FCCP), and rotenone/antimycin A (R/AA) were used at final concentrations of 0.5, 1, and 0.5µM respectively. Mitochondrial stress test parameters were calculated as described previously^28^.

### Glycolytic rate assay

Cells were seeded as described above with 10µM Y-27632. Compounds for the glycolytic rate assay (Agilent, cat. no. 103344-100) were reconstituted in XF DMEM (Table 1). Final injected concentrations were 0.5µM R/AA and 50mM 2-deoxyglucose (2-DG). 4 reads were permitted between injections. A titration curve of HCl injections was carried out on a per-plate basis to determine the buffer factor of the media according to the manufacturer’s specifications. Extracellular acidification rate (ECAR) data were converted to proton efflux rate (PER) according to the manufacturer’s instructions. Glycolytic rate assay parameters were calculated as follows: mitoPER: *[CO_2_ contribution factor] * [mitoOCR]*, mitoOCR: *[fourth baseline OCR read] – [minimum OCR rate measurement following R/AA injection]*, PER(basal): *fourth baseline PER read*, glycoPER: *[baseline PER] – [mitoPER]*, compensatory glycolysis: *[maximal PER measurement after R/AA] – [basal glycoPER]*, maximal glycolysis: *[maximal PER measurement after R/AA injection] – [post-2DG acidification]*, post-2DG acidification: *minimum rate measurement following 2DG injection*.

### Immunocytochemistry

Astrocyte fate specification was confirmed via immunostaining for the markers GFAP, S100B, and aquaporin-4 (AQP4) (Table 2). For these works, cells were seeded at a density of 5.0×10^4^ cells/cm^2^ onto Geltrex-coated 8-chamber slides (Merck, cat. no. PEZGS0816) in seeding media supplemented with 10µM Y-27632. For Translocase of the Outer Mitochondrial Membrane 20 (TOMM20) and nuclear factor kappa-light-chain-enhancer of activated B cells (NFκB) immunostaining, cells were seeded at a density of 2.0×10^4^ cells per well onto 8-chamber slides in seeding media (for NFκB staining cells were supplemented with 10µM Y-27632). The next day, after 2h acclimatisation to assay media, cells were washed once with ice-cold 0.01M PBS and fixed in 4% formaldehyde solution (Merck, cat. no. 1.00496.5000) for 15 minutes at RTP. Wells were rinsed thrice with 0.01M PBS, PBS fully aspirated, and stored at −20°C until staining. Immediately prior to staining, coverslips were warmed to RTP and washed a further three times with 0.01M PBS before blocking in 3% w/v BSA in 0.1% PBS-Triton X100 (Merck, cat. no. X100-500ML) for 15 minutes at RTP. Primary antibody incubation immediately succeeded blocking (all antibodies were diluted in 3% w/v BSA in 0.05% PBS-Tween 20 [Merck, cat. no. P1379-100ML]; for a list of primary antibodies see Table 2) for 15 minutes at RTP before storage at 4°C overnight. The next day, primary antibodies were removed and wells washed thrice with 0.01M PBS prior to 1h RTP incubation with a second primary antibody (if required), washed a further three times with 0.01M PBS, and secondary antibodies were applied for 1h RTP in darkness. Wells were washed, and the nuclear counterstain DAPI (Merck, D9542) applied at 1µg/mL (0.01M PBS) for 15 minutes in darkness. Excess DAPI was removed by washing and the wells allowed to air dry prior to mounting (Invitrogen, P36961). All microscopy was performed using a Zeiss LSM 880 confocal microscope. Images were captured with either a 20x or 63x objective lens using ZEN Black software (version 2.3; version number 14.0.18.201; Zeiss).

**Table 2:** antibodies used for immunocytochemistry.

| Target | Supplier | Catalogue number | Host species | Dilution | RRID |
| --- | --- | --- | --- | --- | --- |
| GFAP | Merck | MAB360 | Mouse | 1/250 | RRID: AB_11212597 |
| S100B | Cell Signalling Technologies | 90393 | Rabbit | 1/250 | RRID: AB_3678716 |
| AQP4 | Invitrogen | PA5-53234 | Rabbit | 1/100 | RRID: AB_2638104 |
| TOMM20 | Abcam | ab56783 | Mouse | 1/100 | RRID: AB_945896 |
| NFkB p65 | Cell Signalling Technology | 6956T | Mouse | 1/100 | RRID: AB_10828935 |
| Mouse (AlexaFluor 488) | Invitrogen | A32723 | Goat | 1/500 | RRID: AB_2633275 |
| Rabbit (AlexFluor 647) | Invitrogen | A21245 | Goat | 1/500 | RRID: AB_2535813 |

### Manual cell counting

Maximum intensity projections were generated from Z-stacks on a per-channel basis in FIJI^30^ with no additional adjustments. Gaussian filters were generated (σ = 2µm) and applied to all images within datasets to remove background fluorescence on a per-channel basis. Image brightness was adjusted on a per-channel basis relative to the least bright image within a dataset. The same parameters were applied to all images within a dataset. Manual cell counting was performed using the Cell Counting module in FIJI^30^ and data were collated in Excel.

### CellProfiler pipelines

CellProfiler^31^ pipelines were used to determine changes to cellular morphology and protein localisation. Raw, unaltered maximum intensity projections were used for each pipeline. Gaussian filters (σ = 100 µm) were used to remove background fluorescence. The *IdentifyPrimaryObjects* module was used to identify nuclei. Cell somas were identified using the *IdentifySecondaryObjects* module in the 488 and 647 channels. Once identified, the *DilateObjects*, *MorphologicalSkeleton* and *MeasureObjectSkeleton* modules were used to determine changes to cellular morphology, and skeletonised images exported using the *SaveImages* module. Other parameters were collected using the *MeasureObjectSizeShape* and *MeasureObjectIntensity* modules. Pixel values were manually converted to µm in Excel using the pixel-to-µm ratio in the image metadata file. Mitochondria were counted using a CellProfiler pipeline adapted from^32^, where oil immersion confocal microscopy (Zeiss LSM 880, 63x objective lens, Zeiss Immersol 518 F, ZEN Black software (version 2.3; version number 14.0.18.201; Zeiss) was used to capture images of single cells. Following identification of nuclei, mitochondria were identified using the *IdentifySecondaryObjects* module and counted. Images were exported from CellProfiler using the *SaveImages* module. Modified images (following background removal and adjustment) were exported using the same module and are displayed in the relevant figures; any adjustments to brightness and contrast for display purposes only were made using FIJI (as described above) and applied equally to all images on a per-channel basis.

### Immunoblotting

Cells were seeded into 6-well plates at a density of 3×10^5^ cells per well and transitioned to assay media as described above. Cells were washed once with ice-cold 0.01M PBS and lysed with lysis buffer (Table 3). Samples were scraped thoroughly on ice, collected, and immediately stored at −80°C for ≥24h. Protein content was estimated via Bradford assay (Bio-Rad, cat. no. 5000006) as described previously^28,29^. Immunoblotting samples were prepared as previously reported^28^, aliquoted and stored at −80°C until use. For SDS-PAGE immunoblotting, 10µg protein was loaded per well into hand-cast 10% (Figure 7) or 15% (Figure 8) polyacrylamide resolving gels (4% stacking gels were used for all samples). Samples were run at 90V for 15 minutes to clear the stacking gel, and resolved by running at 150V for 90 minutes. Gels were wet transferred to nitrocellulose membranes (Bio-Rad, cat. no. 1620146) in 10% methanol tris-glycine buffer at 110V for 70 minutes. A Ponceau stain (Merck, P7170) was used to confirm successful transfer. Ponceau stain was removed by washing with TBS. Membranes were blocked using Odyssey blocking buffer (Li-COR Biosciences, cat. no. 927-60001), 1h RTP. Target antibodies were applied consecutively (Table 4) at 4°C overnight or 1h RTP. Membranes were scanned using a Li-COR Odyssey CLX scanner. Densitometric analysis was performed using ImageStudio Lite v5.2 (Li-COR Biosciences) and fluorescence intensity values exported to Excel. Data are expressed as either [protein of interest]/[loading control], [phosphoprotein band intensity]/[total protein band intensity], or [phosphoprotein band intensity]/[loading control]. Full, uncropped immunoblots can be found in Supplementary Figures 2-7.

**Table 3:**
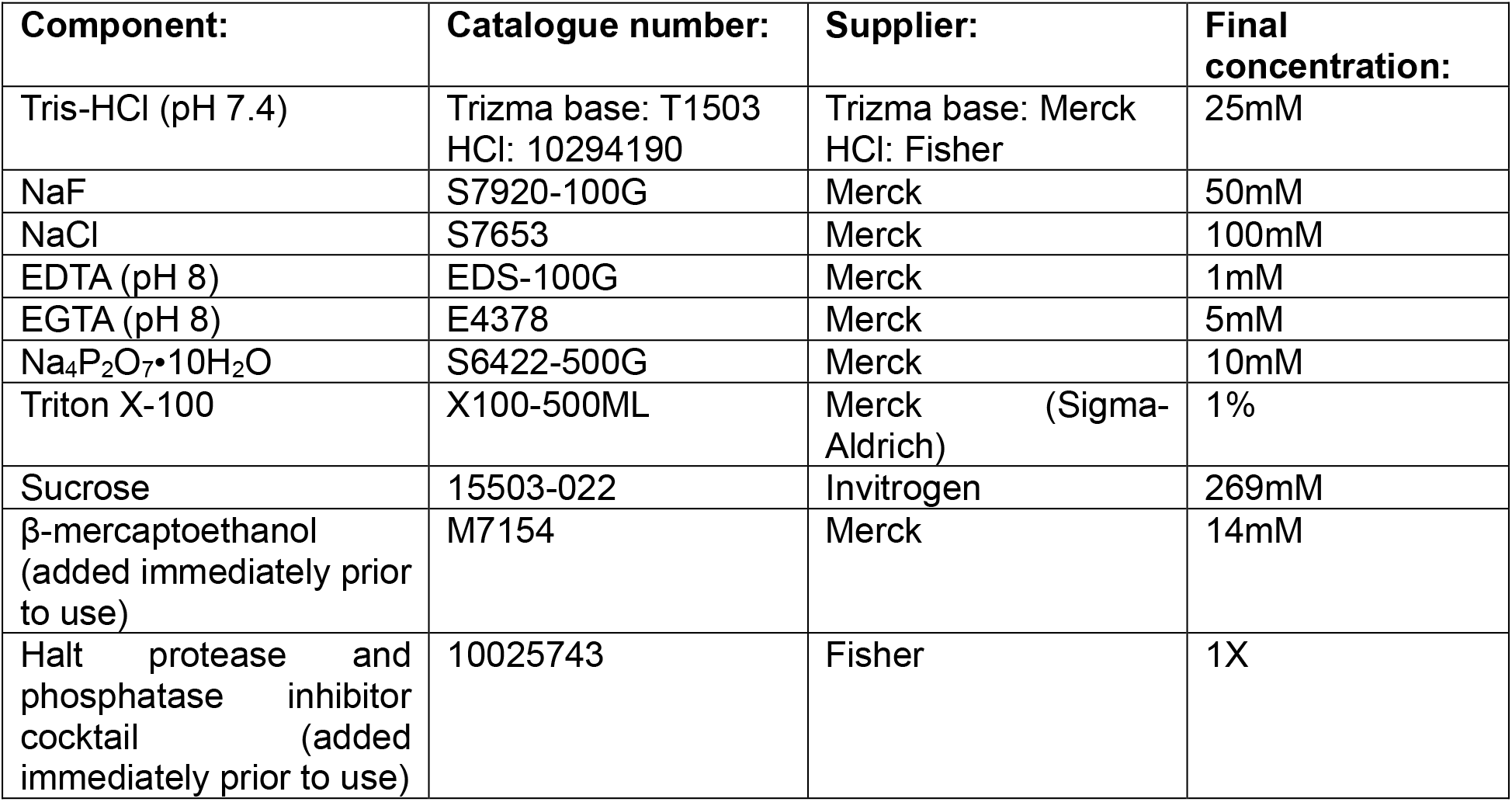
Lysis buffer composition.

| Component: | Catalogue number: | Supplier: | Final concentration: |
| --- | --- | --- | --- |
| Tris-HCl (pH 7.4) | Trizma base: T1503<br>HCl: 10294190 | Trizma base: Merck<br>HCl: Fisher | 25mM |
| NaF | S7920-100G | Merck | 50mM |
| NaCl | S7653 | Merck | 100mM |
| EDTA (pH 8) | EDS-100G | Merck | 1mM |
| EGTA (pH 8) | E4378 | Merck | 5mM |
| $\text{Na}_4\text{P}_2\text{O}_7 \cdot 10\text{H}_2\text{O}$ | S6422-500G | Merck | 10mM |
| Triton X-100 | X100-500ML | Merck (Sigma-Aldrich) | 1% |
| Sucrose | 15503-022 | Invitrogen | 269mM |
| β-mercaptoethanol<br>(added immediately prior to use) | M7154 | Merck | 14mM |
| Halt protease and phosphatase inhibitor cocktail<br>(added immediately prior to use) | 10025743 | Fisher | 1X |

**Table 4:**
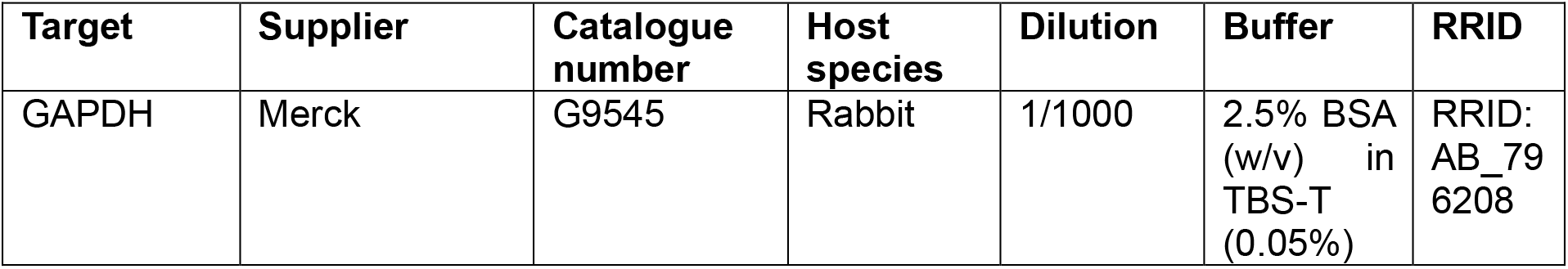
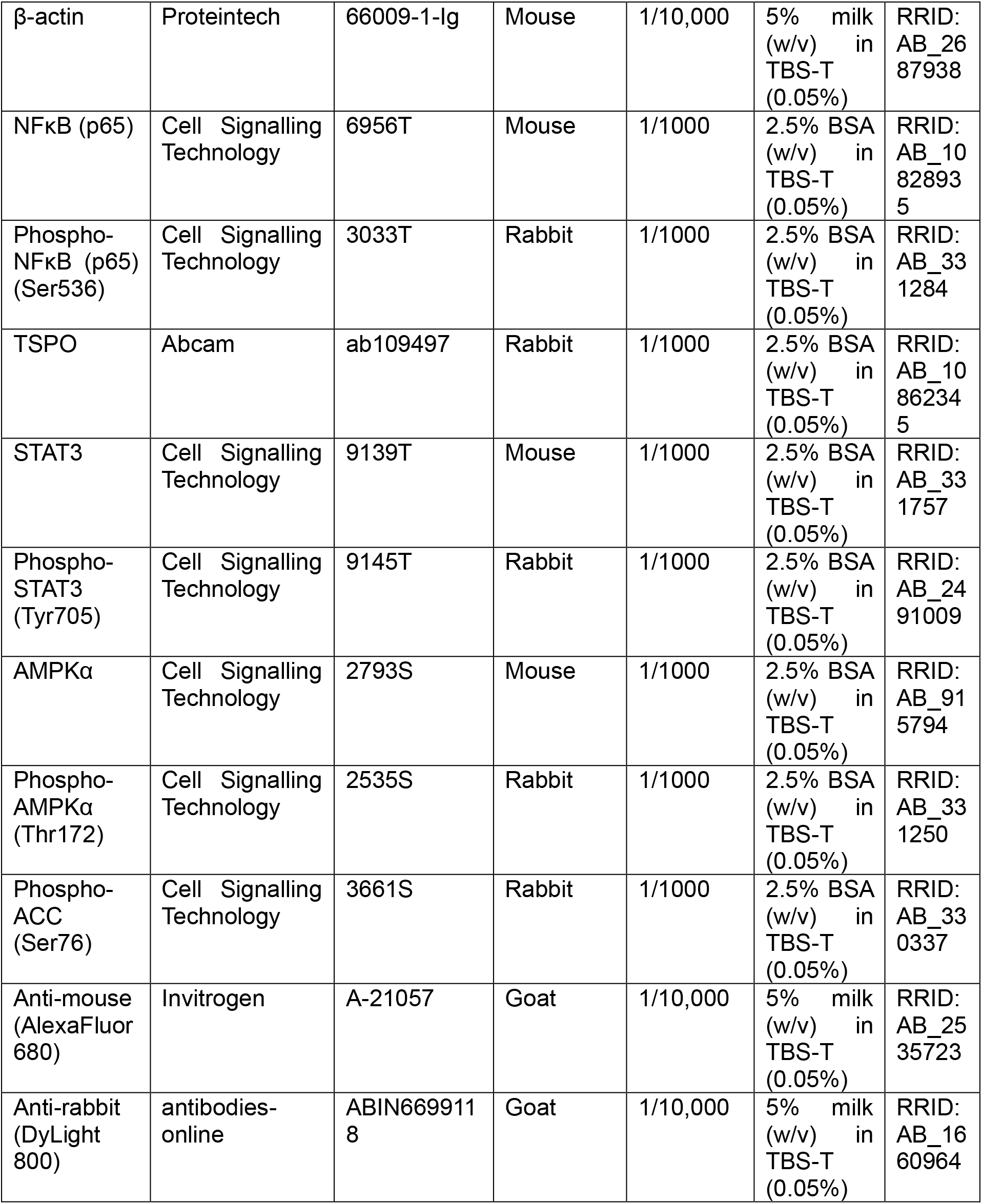
antibodies for immunoblotting.

### Cytokine release profiling

A Human Cytokine Array (Bio-Techne, cat. no. ary022b) was used to probe for changes to the astrocyte secretome in LHON according to the manufacturer’s specifications. Briefly, conditioned media (∼18-24h conditioning time) samples were collected and centrifuged at 1000*xg*, 4°C, for 10 minutes. The supernatant was retained, aliquoted, and stored at −80°C until analysis. Equal volumes of conditioned media from 3 independent replicates (totalling 1.5mL per group) were applied per membrane. Membranes were scanned using a Licor Odyssey CLX scanner (Li-COR Biosciences). ImageStudio was used for densitometric analysis and results exported to Excel for further processing. Background fluorescence from negative control spots was subtracted from all values per membrane. Data were then thresholded to 2.0×10^5^ RFU; any datapoint that achieved this threshold across any of the phenotypes was taken forward for further processing. Data that did not reach this threshold in any single group were not processed further. Data were then expressed as log_2_ fold change from control; all data meeting these criteria are presented in Figure 9. Complete array data are presented in Supplementary Figure 8.

### Data handling and statistical analysis

For the purposes of this study, one replicate was defined as a distinct well, chamber, or dish; as these data were generated from donor-derived stem cells, a true biological replicate (a distinct organism) cannot be generated. Automated image analyses count all cells within a given field of view. Thus, in imaging studies, fields of view were regarded as technical replicates, and are displayed in imaging data distinguished by shape and colour. For imaging studies, we therefore calculated mean values per well, with the well being treated as the biological replicate. Biological replicates are displayed in figures pertaining to imaging studies as solid shapes. Statistical analyses were performed only on biological replicates to avoid oversampling. Extracellular flux analysis data were recorded and normalised within Wave software (version 2.6.4.24, Agilent Technologies) and exported to Excel for parameter calculations. Raw data were processed using Excel and statistical analyses were performed using GraphPad Prism for Windows (version 11.0.1; Graphpad Software, Boston, Massachusetts, USA). Prior to statistical analysis, all data underwent outlier detection and data normality were determined as described previously^28^ with minor modifications: additional normality tests were run (Anderson-Darling test, Shapiro-Wilk test; α=0.05 in all instances). Statistical tests were applied according to the normality of the datasets as determined by normality testing. The threshold for statistical significance was set as p=0.05. Data are displayed as mean±standard error of the mean.

## Results

Having generated and validated astrocytes through the expression of the canonical *in vitro* identity markers GFAP and S100B (Supplementary Figure 1), we next profiled the effect of m.14484T>C-LHON on astrocyte GFAP and S100B expression to test the hypothesis that LHON would upregulate intracellular levels of these proteins.

### LHON-affected astrocytes showed reduced intracellular GFAP and S100B expression

Like the healthy control astrocytes, LHON-affected astrocytes were immunopositive for GFAP and S100B (Figure 1A), though the phenotype of the cells was affected by LHON status. The number of nuclei differed but we were unable to detect where the difference lay (Figure 1B). Nuclear fluorescence intensity was significantly increased in both LHON phenotypes (Figure 1C). There was a significant increase in nuclear area (Figure 1D), though post-hoc testing did not reveal where the difference lay. There was no significant difference in nuclear length (Figure 1E). Although cells across all phenotypes were GFAP-and S100B-expressing (Figure 1F,H), the intracellular intensities of these proteins were significantly reduced relative to the healthy control line (Figure 1G,I), though no significant difference in GFAP (p=0.1564) or S100B (p=0.5391) intensity between LHON phenotypes was detected. Concordantly, both phenotypes displayed a significant reduction in GFAP-and S100B-positive areas (Figure 1J,K), with the R phenotype showing significantly reduced intracellular GFAP in comparison to the NR phenotype (Figure 1J). We also noted the apparent presence of intra-and extra-cytoplasmic inclusions of GFAP in images of the LHON-affected cells, which appeared to increase relative to the unrelated healthy control (Figure 1L; magenta arrows: intracytoplasmic inclusions, yellow arrows: extracytoplasmic inclusions); these may be linked to inappropriate GFAP cleavage^33^ or leakage^34^.

**Figure 1:**
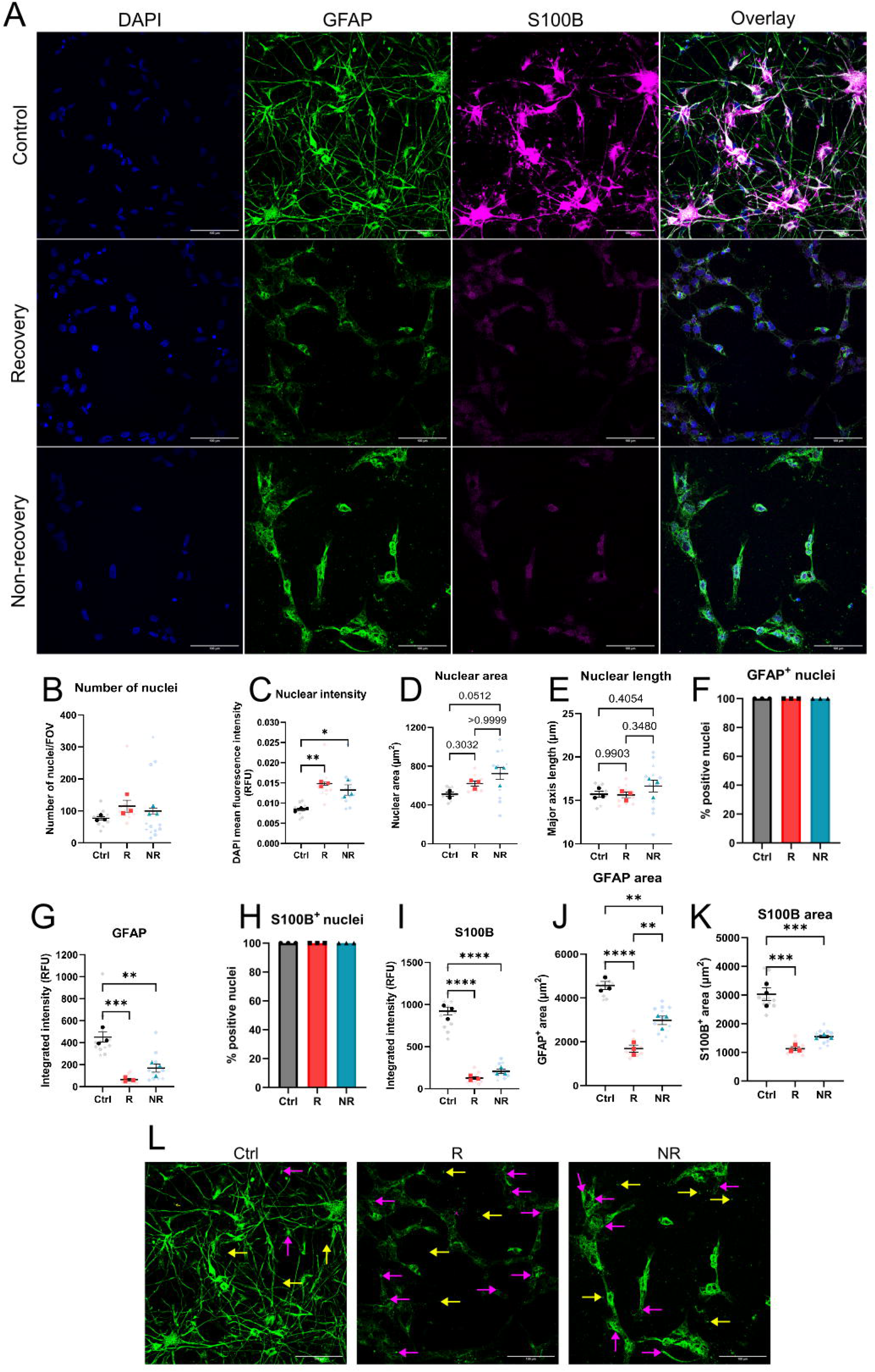
LHON disrupts astrocyte morphology and reduces intracellular GFAP and S100B expression. A: Immunocytochemical staining of LHON-affected astrocytes compared to healthy control. Blue: DAPI, green: GFAP, magenta: S100B. B: Number of nuclei per FOV. There were more nuclei per FOV in LHON (p_Kruskal-Wallis_=0.025, Kruskal-Wallis statistic=5.956, p_Ctrl-R_=0.0512, p_Ctrl-NR_=0.3032, p_R-NR_>0.99). C: nuclear intensity of DAPI was significantly increased in LHON (p_ANOVA_=0.0030, F=17.87, p_Ctrl-R_=0.0029, p_Ctrl-NR_=0.0117, p_R-NR_=0.3946). D: LHON increased nuclear area (p_Kruskal-Wallis_=0.025, Kruskal-Wallis statistic=5.596, p_Ctrl-R_=0.3032, p_Crtl-NR_=0.0512, p_R-NR_>0.99). E: LHON did not affect nuclear length (p_ANOVA_=0.3140, F_2,6_=1.414). F: 100% of nuclei associated with GFAP positivity. G: LHON reduced GFAP integrated intensity (p_ANOVA_=0.0005, F=34.49, p_Ctrl-R_=0.0005, p_Ctrl-NR_=0.0026, p_R-NR_=0.1564). H: 100% of nuclei associated with S100B positivity. I: LHON reduced S100B integrated intensity (p_Kruskal-Wallis_=0.0036, Kruskal-Wallis statistic=7.2, p_Ctrl-R_=0.0219, p_Ctrl-NR_=0.5391, p_R-NR_=0.5391). J: LHON reduced GFAP-positive cell area (p_ANOVA_<0.0001, F=62.87, p_Ctrl-R_<0.0001, p_Ctrl-NR_=0.0020, p_R-NR_=0.0058). K: LHON reduced S100B-positive cell area (S100B p_ANOVA_=0.0001, F=53.6, p_Ctrl-R_=0.0002, p_Ctrl-NR_=0.0006, p_R-NR_=0.1516). L: Duplicates of panel A with annotations indicating putative GFAP inclusions. LHON was associated with an increased number of intracytoplasmic (yellow arrows) and extracytoplasmic (magenta arrows) GFAP puncta. n=3 (solid dots; B-K), 4 FOV per replicate (transparent shapes; B-E, G, I, J, K). Data are mean±SEM. *p<0.05, **p<0.01, ***p<0.001, ****p<0.0001. Ordinary one-way ANOVA with Tukey’s multiple comparisons test (C, E, G, J, K) or Kruskal-Wallis test with Dunn’s multiple comparisons (B, D, I). DAPI: 4′,6-Diamidino-2-phenylindole, GFAP: glial fibrillary acidic protein, S100B: S100 calcium binding protein B, FOV: field(s) of view, Ctrl: healthy control, R: m.14484T>C-LHON with spontaneous visual recovery, NR: m.14484T>C-LHON without spontaneous visual recovery, RFU: relative fluorescence units. Scale bars: 100µm.

### LHON reduced astrocyte morphological complexity

Having identified that LHON reduced intracellular GFAP and S100B expression in our iPSC-derived astrocytes, we next sought to understand the relationship between LHON and astrocyte morphological complexity. Changes to astrocyte morphological complexity are noted in many pathophysiological contexts^19,35^ and are classically considered to consist of somatic enlargement, hypertrophy, and increased GFAP expression^35^; though this is not always the case^23,35^. Thus, the morphological complexity of LHON-affected astrocytes was interrogated. Identified nuclei were used as ‘seeds’ from which cell networks were automatically constructed using GFAP-expressing areas (as areas of GFAP expression appeared to be greater than S100B expression across phenotypes, Figure 1A). Skeletonised constructions were divided into trunks (basic processes extended from cell somas), branches (points where trunks divide), and termini (the number of branch endpoints) (Figure 2A,B). There was no difference in the number of basic processes projected per cell (Figure 2C), however the number of branches was significantly reduced in both LHON phenotypes relative to control (Figure 2D), suggesting that LHON did indeed reduce astrocyte morphological complexity. Similarly, the number of termini was significantly reduced in LHON-R astrocytes (Figure 2E). Seeking to further quantify the extent of these changes, the total length of all processes across the LHON phenotypes was examined. LHON reduced the length of processes relative to healthy control (Figure 2F), with a significant difference in process length between LHON phenotypes (p_R-NR_=0.0138).

**Figure 2:**
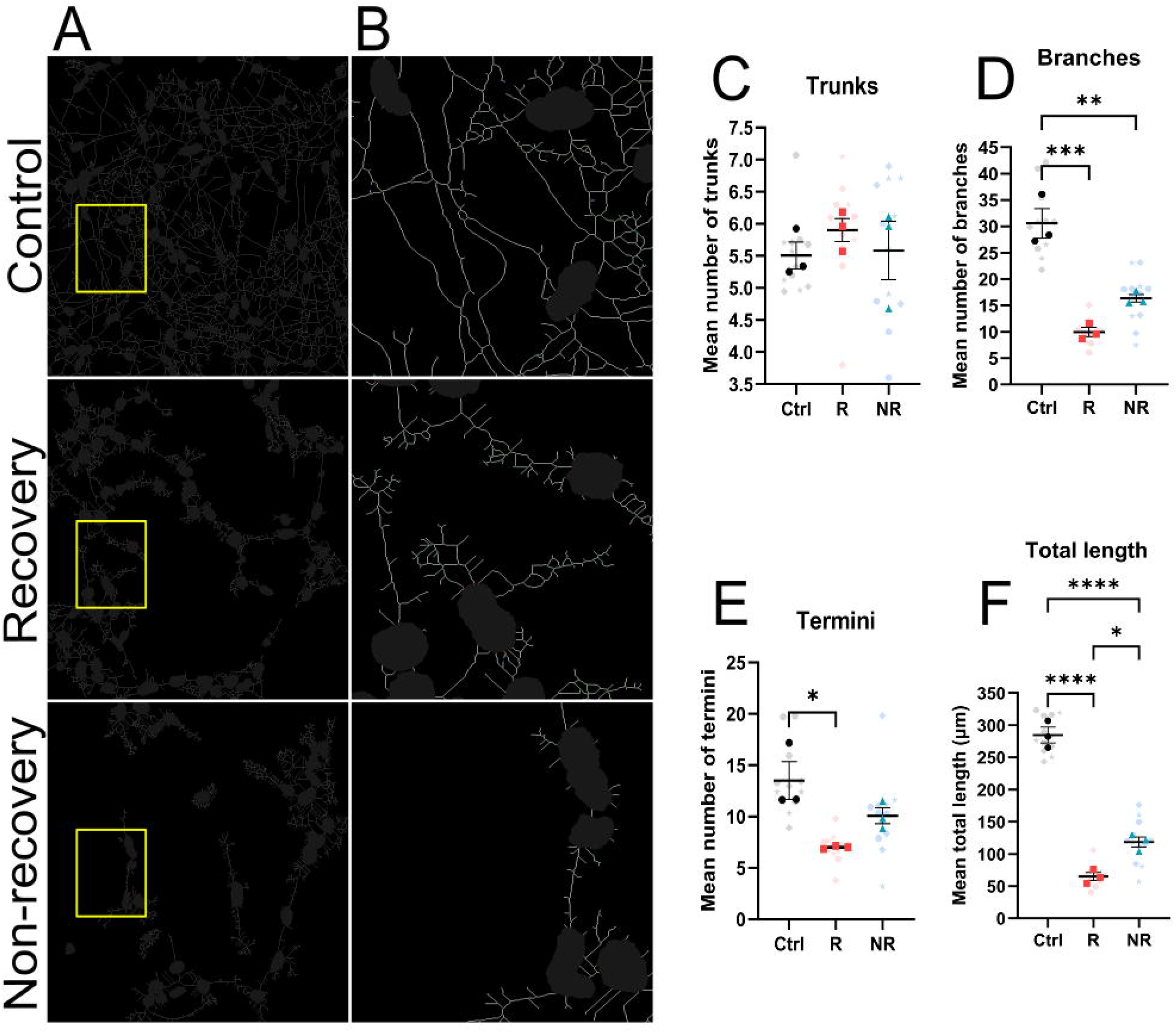
LHON reduces astrocyte morphological complexity. A: skeletonised images of GFAP-positive somas from Figure 1A. B: ROIs, denoted by yellow box in A. C: LHON did not affect the number of trunks projected per cell (p_ANOVA_=0.6458, F=0.4707). D: LHON significantly reduced the number of branches per cell (p_ANOVA_=0.0004, F=36.95, p_Ctrl-R_=0.0004, p_Ctrl-NR_=0.0028, p_R-NR_=0.0883). E: LHON significantly reduced the number of termini per cell (p_Kruskal-Wallis_=0.0036, Kruskal-Wallis statistic=7.2, p_Ctrl-R_=0.0219, p_Ctrl-NR_=0.5391, p_R-NR_=0.5391). F: LHON significantly reduced the total length of processes (p_ANOVA_<0.0001, F=159.8, p_Ctrl-R_<0.0001, p_Ctrl-NR_<0.0001, p_R-NR_=0.0138). n=3 (solid dots; B-F), 4 FOV per replicate (transparent shapes; B-F). Data are mean±SEM. *p<0.05, **p<0.01, ***p<0.001, ****p<0.0001. Ordinary one-way ANOVA with Tukey’s multiple comparisons test (C, D, F) or Kruskal-Wallis test with Dunn’s multiple comparisons (E). LHON: Leber’s Hereditary Optic Neuropathy, Ctrl: healthy control, R: m.14484T>C-LHON with spontaneous visual recovery, NR: m.14484T>C-LHON without spontaneous visual recovery, ROI: region of interest.

### LHON-NR astrocytes showed AQP4 mislocalisation

AQP4 is a key player in CNS water transport expressed on astrocyte endfeet and modulated by pathological status^36,37^. To understand if the latter was the basis of reduced astrocyte morphological complexity, we probed for aquaporin-4 expression. Co-staining for GFAP and AQP4 (Figure 3A) confirmed that astrocytes of all phenotypes expressed AQP4 and again confirmed a reduction in GFAP area and intracellular intensity in LHON (Figure 3A-C). We next interrogated the possibility that LHON may alter GFAP subcellular localisation, following on from our observation of GFAP inclusions in LHON-affected astrocytes (Figure 1L). Nuclear intensity of GFAP was significantly reduced in LHON relative to control, specifically the R but not NR phenotype (Figure 3D), while GFAP intensity at cell edges was significantly reduced in both LHON phenotypes (Figure 3E). Edge:nucleus GFAP intensity was also reduced (Figure 3F), reinforcing our results.

**Figure 3:**
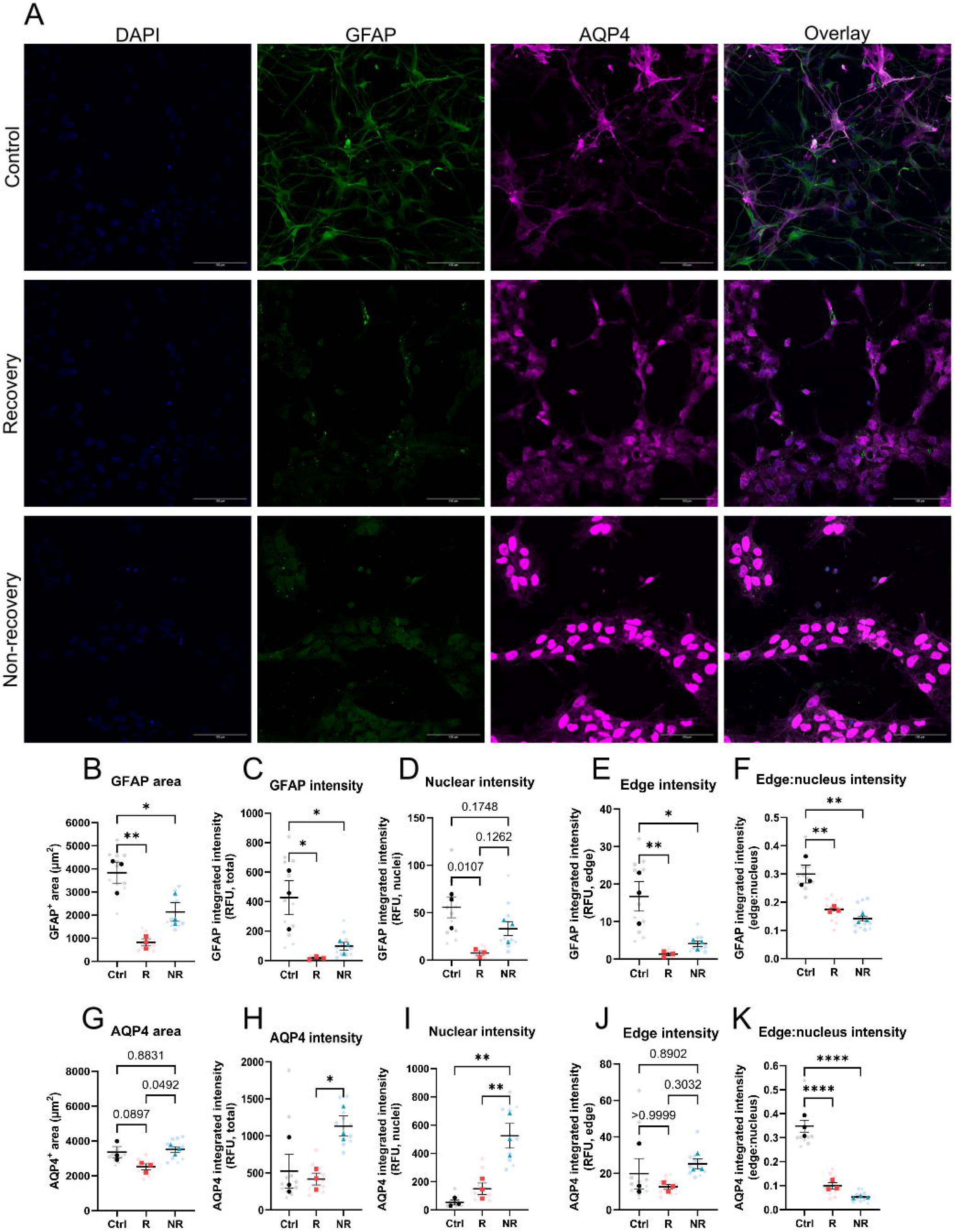
LHON induces AQP4 mislocalisation in astrocytes. A: immunocytochemical staining of LHON-affected astrocytes in comparison to healthy control. Blue: DAPI, green: GFAP, magenta: AQP4. B: GFAP^+^ area was significantly reduced in LHON affected astrocytes (p_ANOVA_=0.0031, F=17.62, p_Ctrl-R_=0.0025, p_Ctrl-NR_=0.0364, p_R-NR_=0.0909). C: LHON significantly reduced GFAP intracellular intensity (p_ANOVA_=0.0124, F=9.972, p_Ctrl-R_=0.0132, p_Ctrl-NR_=0.0341, p_R-NR_=0.6981). D: LHON reduced nuclear intensity of GFAP (p_ANOVA_=0.0131, F=9.741, p_Ctrl-R_=0.0107, p_Ctrl-NR_=0.1748, p_R-NR_=0.1262). E: GFAP edge intensity was significantly reduced in LHON-affected astrocytes (p_ANOVA_=0.0075, F=12.33, p_Ctrl-R_=0.0083, p_Ctrl-NR_=0.0206, p_R-NR_=0.6915). F: edge:nucleus GFAP intensity was significantly reduced in LHON (p_ANOVA_=0.0027, F=18.58, p_Ctrl-R_=0.0089, p_Ctrl-NR_=0.0029, p_R-NR_=0.5063). G: LHON significantly reduced AQP4^+^ areas (p_ANOVA_=0.0440, F=5.497, p_Ctrl-R_=0.0897, p_Ctrl-NR_=0.8831, p_R-NR_=0.0492). H: Intracellular AQP4 intensity was increased in LHON-affected astrocytes (p_ANOVA_=0.0412, F=5.684, p_Ctrl-R_=0.8912, p_Ctrl-NR_=0.0831, p_R-NR_=0.0467). I: LHON increased nuclear AQP4 intensity (p_ANOVA_=0.0024, F=19.38, p_Ctrl-R_=0.4936, p_Ctrl-N_=0.0026, p_R-NR_=0.0080). J: AQP4 intensity at cell edges was unchanged by LHON (p_Kruskal-Wallis_=0.2964, Kruskal-Wallis statistic=2.756). K: edge:nucleus intensity ratio of AQP4 was significantly reduced in LHON-affected cells (p_ANOVA_<0.0001, F=95.58, p_Ctrl-R_<0.0001, p_Ctrl-NR_<0.0001, p_R-NR_=0.1827). n=3 (solid dots; B-K), 4 FOV per replicate (transparent shapes; B-K). Data are mean±SEM. *p<0.05, **p<0.01, ****p<0.0001. Ordinary one-way ANOVA with Tukey’s multiple comparisons test (B-I, K) or Kruskal-Wallis test with Dunn’s multiple comparisons (J). DAPI: 4′,6-Diamidino-2-phenylindole, GFAP: glial fibrillary acidic protein, AQP4: aquaporin-4, FOV: field(s) of view, Ctrl: healthy control, R: m.14484T>C-LHON with spontaneous visual recovery, NR: m.14484T>C-LHON without spontaneous visual recovery, RFU: relative fluorescence units. Scale bars: 100µm.

Consistent with this, AQP4-positive cell areas were significantly reduced in LHON (Figure 3G); post-hoc testing revealed that the difference lay between LHON-affected groups (p_Ctrl-R_=0.0897, p_Ctrl-NR_=0.8831, p_R-NR_=0.0492). Likewise, total AQP4 intensity was significantly increased in m.14484T>C-LHON-NR affected astrocytes relative to m.14484T>C-LHON-R astrocytes (Figure 3H). Further analysis revealed LHON phenotype-dependent effects on AQP4 localisation: nuclear intensity of AQP4 was significantly increased in the NR phenotype relative to both healthy control and the R phenotype (Figure 3I). Whilst there was no change in AQP4 intensity at cell edges (Figure 3J), edge:nucleus intensity of AQP4 was significantly reduced in both LHON phenotypes (Figure 3K) suggesting that m.14484T>C-LHON modulates AQP4 subcellular localisation in a manner potentially attributable to impediments in protein translocation.

### LHON did not affect astrocyte mitochondrial load

Hypothesising that our prior findings were attributable to the Complex I dysfunction associated with LHON mutations^9,38,39^, and further hypothesising that this would be attributable to impaired mitophagy:mitobiogenesis in LHON-affected cells^40^, mitochondrial load was evaluated via immunostaining of astrocytes for Translocase of the Outer Mitochondrial Membrane 20 (TOMM20; Figure 4A). By adapting an existing pipeline^32^, mitochondrial load per cell was determined (Figure 4B). No significant difference in mitochondrial number per cell was identified, though the R phenotype trended towards a reduction (Figure 4B). As a secondary measure of mitochondrial load, mean fluorescence intensity of TOMM20 immunostaining per cell was quantified and also determined no significant difference between any of our groups (Figure 4C).

**Figure 4:**
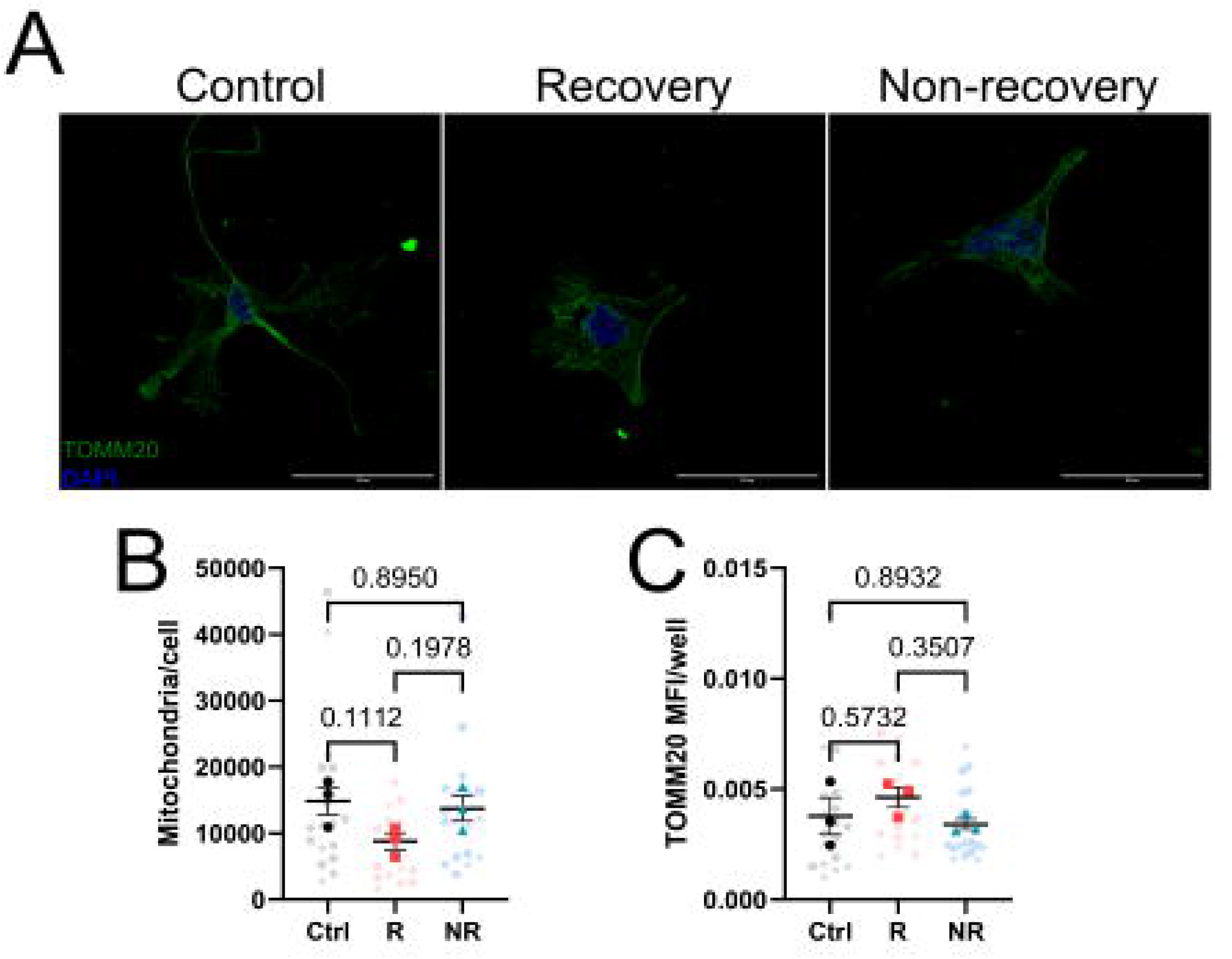
LHON did not affect astrocyte mitochondrial load. A: immunocytochemical staining of TOMM20 and DAPI in astrocytes. B: LHON did not affect the number of mitochondria per cell (p_ANOVA_=0.1055, F_2,6_=3.349). C: There was no effect of LHON on TOMM20 MFI (p_ANOVA_=0.3639, F=1.202). n=3 (solid dots; B,C), 6 FOV per replicate (transparent shapes; B,C). Data are mean±SEM. Ordinary one-way ANOVA with Tukey’s multiple comparisons test (B,C). DAPI: 4′,6-Diamidino-2-phenylindole, TOMM20: translocase of the outer mitochondrial membrane 20, FOV: field(s) of view, Ctrl: healthy control, R: m.14484T>C-LHON with spontaneous visual recovery, NR: m.14484T>C-LHON without spontaneous visual recovery, MFI: mean fluorescence intensity. Scale bars: 50µm.

### LHON reduced mitochondrial metabolism with differences between phenotypes

Our findings thus far indicated that the mitochondrial DNA mutation associated with m.14484T>C-LHON was sufficient to drive a reduction in astrocyte morphological complexity, reductions in intracellular GFAP and S100B, and an overall increase in AQP4 expression with mislocalisation of AQP4 to the nucleus evident in the NR phenotype, but that this could not be attributed to modulation of mitochondrial load as determined using our methodologies.

An alternative explanation for our observations was that the LHON-R and -NR phenotypes may be associated with differentially modulated mitochondrial function. We tested this hypothesis via live cell extracellular metabolic flux analysis (Figure 5). The mitochondrial stress test (Figure 5A) was selected to assay mitochondrial metabolism and stress responses. We found that LHON-NR astrocytes showed a significant reduction in non-mitochondrial respiration, i.e., non-mitochondrial O_2_ consumption, relative to healthy control (Figure 5B). Concurrently, these cells showed a deficit in basal (Figure 5C) and maximal (Figure 5D) mitochondrial respiration relative to both the R and healthy control phenotype; though the NR phenotype showed no significant change in H^+^ leak, a factor which was significantly upregulated in the R phenotype (Figure 5E). Both LHON phenotypes showed statistically significant reductions in ATP-linked OCR relative to healthy control and each other (Figure 5F), suggestive of impaired coupling efficiency which mirrored this trend (Figure 5G). Spare capacity was significantly upregulated in both LHON phenotypes (Figure 5H), perhaps due to the calculation used for this parameter (see Methods). We suggest that this may be explained by our observation of reduced basal metabolism allowing for a transiently greater increase in mitochondrial respiration in response to FCCP administration (Figure 5A) that is not maintained long-term in contrast to the healthy control.

**Figure 5:**
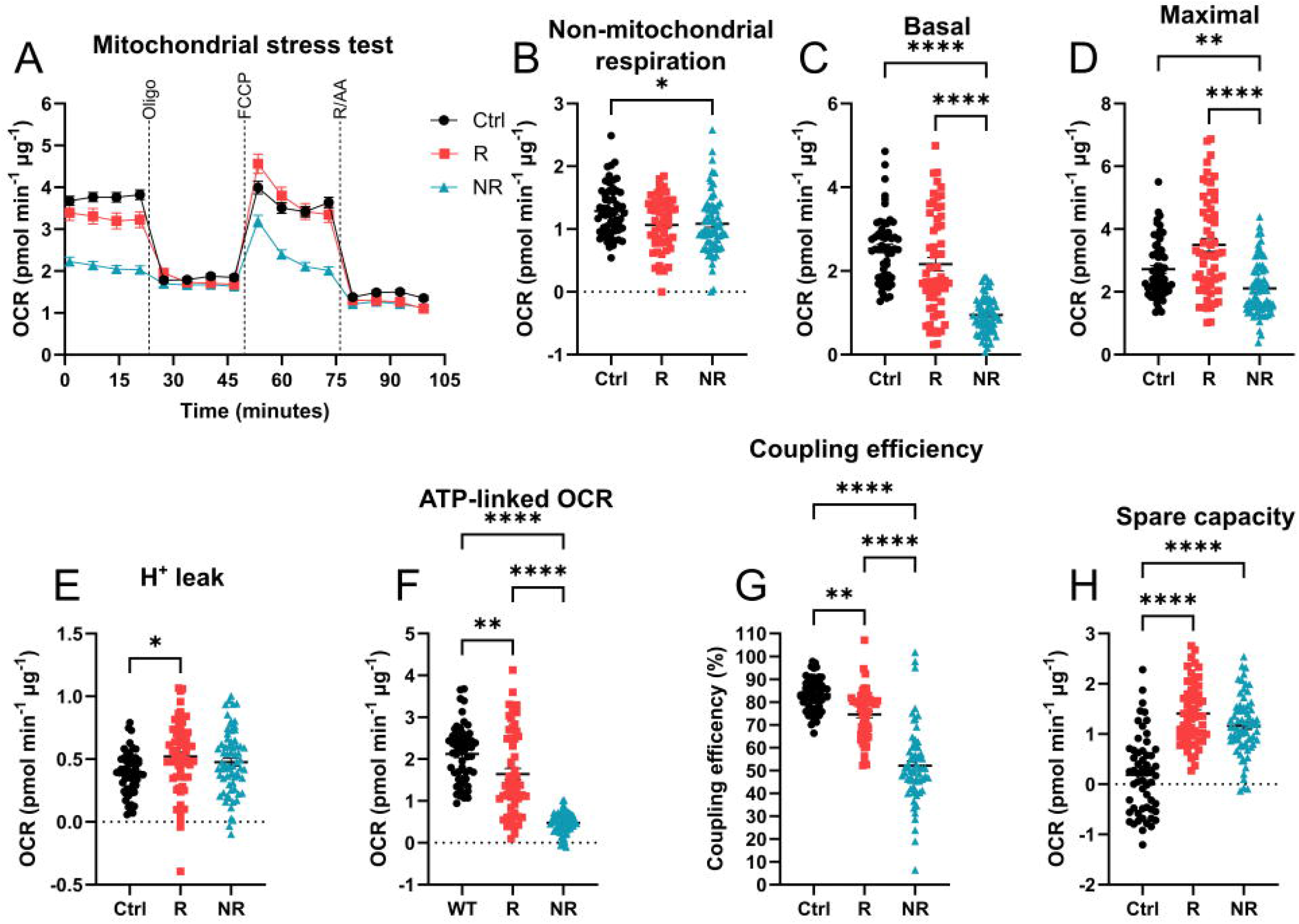
LHON reduced astrocyte mitochondrial metabolism. A: schematic showing the mitochondrial stress test. B: LHON significantly reduced non-mitochondrial respiration (p_Kruskal-Wallis_=0.0122, Kruskal-Wallis statistic=8.806, p_Ctrl-R_=0.0542, p_Ctrl-NR_=0.0170, p_R-NR_>0.9999). C: LHON significantly reduced basal mitochondrial respiration in astrocytes (p_ANOVA_<0.0001, F=60.19, p_Ctrl-R_=0.0553, p_Ctrl-NR_<0.0001, p_R-NR_<0.0001). D: LHON significantly reduced astrocyte maximal respiration (p_Kruskal-Wallis_<0.0001, Kruskal-Wallis statistic=33.12, p_Ctrl-R_=0.0850, p_Ctrl-NR_=0.0021, p_R-NR_<0.0001). E: H^+^ leak was significantly increased in LHON (p_ANOVA_=0.0136, F=4.401, p_Ctrl-R_=0.0112, p_Ctrl-NR_=0.11, p_R-NR_=0.5552). F: LHON significantly reduced astrocyte ATP-linked OCR (p_ANOVA_<0.0001, F=89.28, p_Ctrl-R_=0.0011, p_Ctrl-NR_<0.0001, p_R-NR_<0.0001). G: Coupling efficiency was reduced in LHON-affected astrocytes (p_Kruskal-Wallis_<0.0001, Kruskal-Wallis statistic=95.7, p_Ctrl-R_=0.0019, p_Ctrl-NR_<0.0001, p_R-NR_<0.0001). H: Spare capacity was significantly increased in LHON-affected astrocytes (p_ANOVA_<0.0001, F=55.03, p_Ctrl-R_<0.0001, p_Ctrl-NR_<0.0001, p_R-NR_=0.0998). n=57-62 wells, data are pooled from across 2 independent plates. Data are expressed as mean±SEM. Ordinary one-way ANOVA with Tukey’s multiple comparisons test (C, E, F, H) or Kruskal-Wallis test with Dunn’s multiple comparisons (B, D, G). *p<0.05, **p<0.01, ***p<0.001, ****p<0.0001. OCR: oxygen consumption rate, Oligo: oligomycin (0.5µM), FCCP: carbonyl cyanide-p-trifluoromethoxyphenylhydrazone (1 µM), R/AA: rotenone/antimycin A mix (0.5µM), Ctrl: healthy control, R: m.14484T>C-LHON with spontaneous visual recovery, NR: m.14484T>C-LHON without spontaneous visual recovery.

### LHON increased astrocyte glycolysis

Having established that LHON drives phenotype-dependent modulation of astrocyte mitochondrial respiration, we next tested the hypothesis that the downregulation of mitochondrial respiration would be concomitant with a compensatory upregulation of glycolysis. We again employed live cell extracellular metabolic flux analysis and assayed glycolytic metabolism using the glycolytic rate assay (Figure 6A). As predicted, LHON-affected astrocytes of both phenotypes showed a significant upregulation of basal proton efflux rate (PER; Figure 6B) and glycoPER (PER attributable to glycolysis; Figure 6C,D) with LHON-NR astrocytes showing a significantly greater increase in glycolysis in comparison to the R phenotype. Compensatory (Figure 6E) and maximal (Figure 6F) glycolysis were likewise significantly upregulated but the difference between LHON phenotypes was maintained only in maximal glycolysis (Figure 6F, p_R-NR_<0.0001). Both LHON phenotypes showed a statistically significant increase in post-2DG acidification relative to the healthy control (Figure 6H). In line with our earlier data (Figure 5), the proportion of mitoOCR/glycoPER was significantly reduced in both LHON phenotypes (Figure 6G), further suggesting that the upregulation of glycolysis was concomitant with a reduction in mitochondrial respiration.

**Figure 6:**
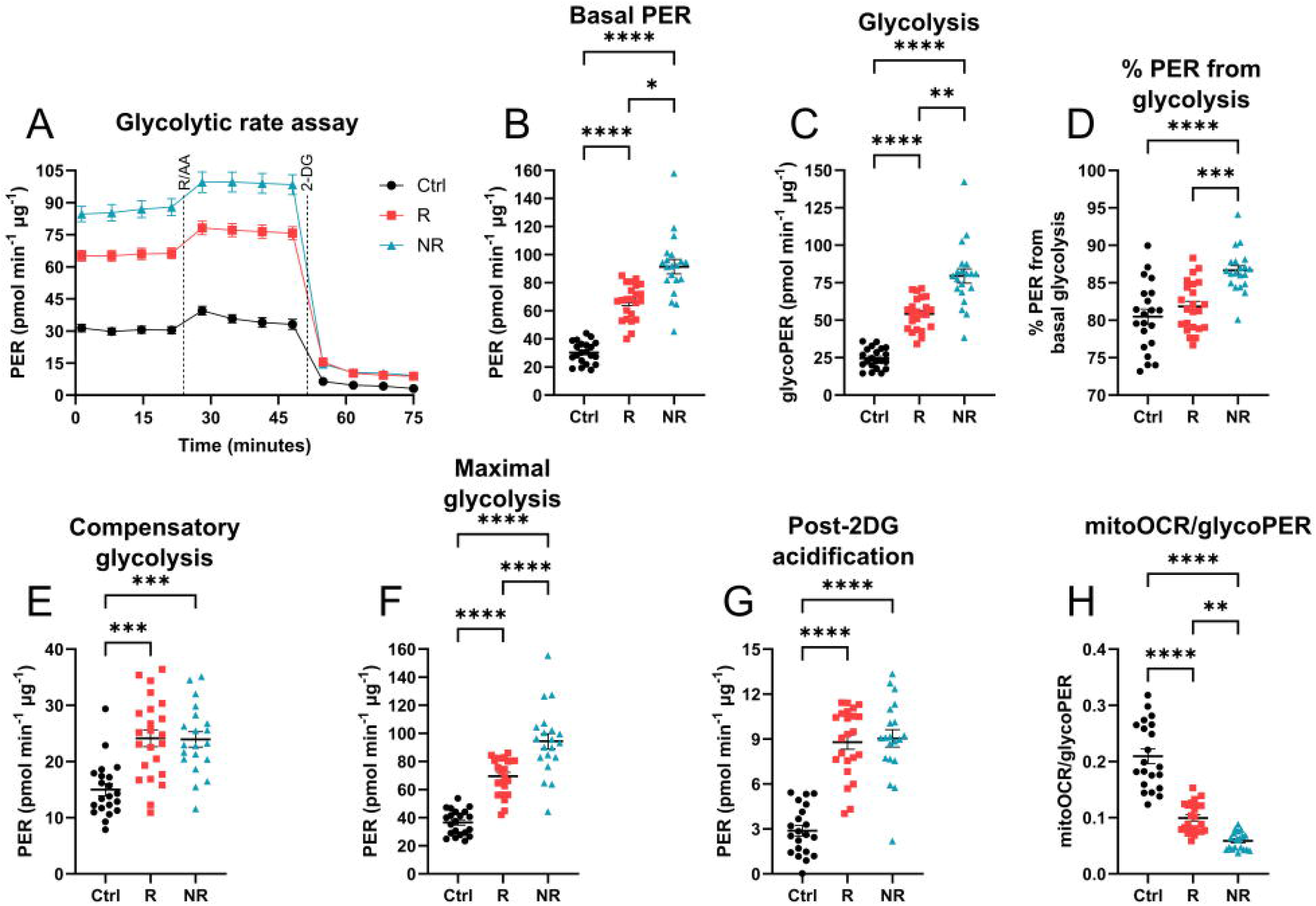
LHON increased astrocyte glycolysis. A: schematic showing the glycolytic rate assay. B: basal PER was significantly increased in LHON-affected astrocytes (p_Kruskal-Wallis_<0.0001, Kruskal-Wallis statistic=43.77, p_Ctrl-R_<0.0001, p_Ctrl-NR_<0.0001, p_R-NR_=0.0171). C: glycolysis-attributable PER was significantly increased by LHON and differed significantly between LHON phenotypes (p_Kruskal-Wallis_<0.0001, Kruskal-Wallis statistic=49.94, p_Ctrl-R_<0.0001, p_Ctrl-NR_<0.0001, p_R-NR_=0.0099). D: percentage of PER from glycolysis was significantly increased in LHON-NR but not R, relative to healthy control (p_ANOVA_<0.0001, F=16.07 p_Ctrl-R_=0.4419, p_Ctrl-NR_<0.0001, p_R-NR_=0.0002). E: compensatory glycolysis was significantly upregulated in both LHON phenotypes (p_Kruskal-Wallis_<0.0001, Kruskal-Wallis statistic=21.8, p_Ctrl-R_=0.0001, p_Ctrl-NR_=0.0002, p_R-NR_>0.9999). F: Maximal glycolysis was significantly increased in LHON-affected astrocytes and different by recovery status (p_ANOVA_<0.0001, F=62.90, p_Ctrl-R_<0.0001, p_Ctrl-NR_<0.0001, p_R-NR_<0.0001). G: Post-2DG acidification was significantly increased in LHON-affected astrocytes but did not differ between LHON phenotypes (p_ANOVA_<0.0001, F=53.12, p_Ctrl-R_<0.0001, p_Ctrl-NR_<0.0001, p_R-NR_=0.9282). H: mitoOCR/glycoPER was significantly reduced in LHON-affected astrocytes relative to healthy control, and differed between LHON phenotypes (p_ANOVA_<0.0001, F=84.98, p_Ctrl-R_<0.0001, p_Ctrl-NR_<0.0001, p_R-NR_=0.0029). n=19-22 wells, data are pooled from 2 independent plates. Data are expressed as mean±SEM. Ordinary one-way ANOVA with Tukey’s multiple comparisons test (D, F-H) or Kruskal-Wallis test with Dunn’s multiple comparisons (B, C, E). *p<0.05, **p<0.01, ***p<0.001, ****p<0.0001. R/AA: rotenone/antimycin A mix (0.5µM), 2DG: 2-deoxyglucose (50mM). Ctrl: healthy control, R: m.14484T>C-LHON with spontaneous visual recovery, NR: m.14484T>C-LHON without spontaneous visual recovery. OCR: oxygen consumption rate. PER: proton efflux rate.

### LHON increased NFκB, but not STAT3 or AMPK, phosphorylation in astrocytes

Recent trends in astrocyte metabolic research have indicated that, in the context of neuroinflammation or neurodegenerative diseases, the astrocyte bioenergetic signature undergoes a shift towards enhanced glycolysis linked to prolonged, unresolved pro-inflammatory stimulation^19^. We thus hypothesised that our data may be explained by chronic inflammation associated with Complex I dysfunction; a continuous insult which does not allow for progression into the important resolutory phase of the inflammatory response. To test this hypothesis we used immunoblotting to assay phosphorylation of Signal Transducer and Activator of Transcription 3 (STAT3), a key mediator of astrocyte inflammatory responses^41,42^ (Figure 7A-C). We found that there was no change in STAT3 phosphorylation at the Tyr705 residue relative to total STAT3 expression (Figure 7A,B) but that total expression of STAT3 was reduced (Figure 7C). In parallel we tested the hypothesis that the bioenergetic shift in astrocytes would be reflected by increased phosphorylation of AMP-activated protein kinase (AMPK; Figure 7D,E) and its downstream target Acetyl-Coenzyme A Carboxylase (ACC; Figure 7F) and found no significant shift in either AMPK expression or phosphorylation of AMPK and ACC (Figure 7D-F).

**Figure 7:**
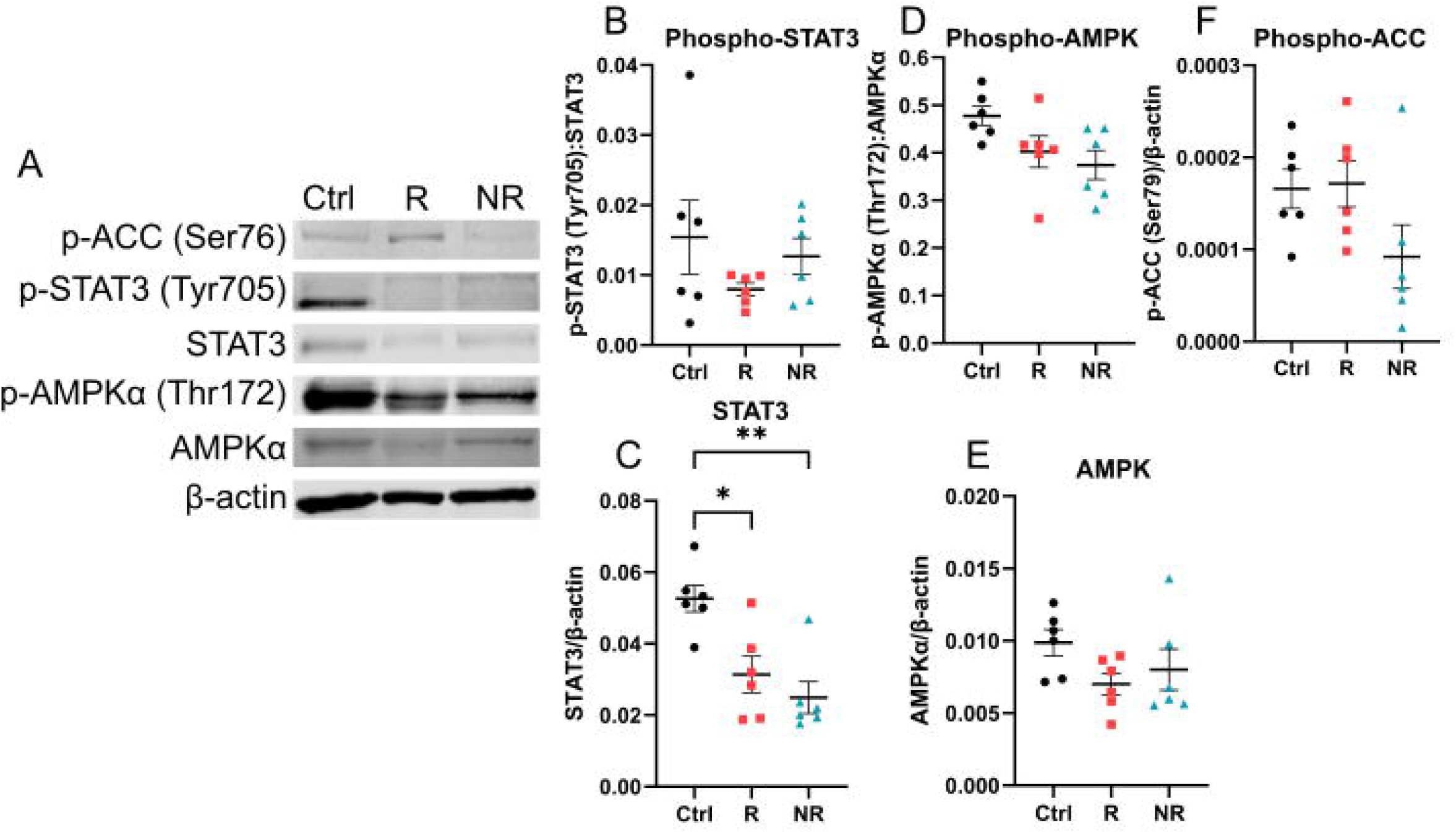
LHON did not affect STAT3, AMPK, or ACC activation in astrocytes. A: representative immunoblots. B: LHON-affected astrocytes showed no significant difference in STAT3 phosphorylation at Tyr705 (p_ANOVA_=0.3264, F=1.207). C: Total STAT3 expression was significantly reduced in LHON affected astrocytes and did not differ between LHON phenotypes (p_ANOVA_=0.0002, F=17.16, p_Ctrl-R_=0.0039, p_Ctrl-NR_=0.0002, p_R-NR_=0.1680). D: AMPK phosphorylation was unaffected by LHON (p_ANOVA_=0.0549, F=3.544). E: Total AMPK expression in astrocytes was unaffected by LHON (p_ANOVA_=0.1870, F=1.879). F: ACC phosphorylation was not affected by LHON (p_ANOVA_=0.11, F=2.566). n=5-6. Data are expressed as mean±SEM. Ordinary one-way ANOVA with Tukey’s multiple comparisons test (B-E). *p<0.05, **p<0.01.

In addition to STAT3, astrocyte inflammatory responses are also governed by nuclear factor kappa-light-chain-enhancer of activated B cells (NFκB) which has been shown to modulate astrocyte immunometabolism^29^. Seeking to explain our finding that LHON-affected astrocytes showed no significant change in STAT3 phosphorylation, we next assayed NFκB activation via immunoblotting. We found a significant upregulation of NFκB phosphorylation in LHON-affected astrocytes (Figure 8A,B). Curiously, total levels of NFκB were significantly reduced (Figure 8C), mirroring our STAT3 results. We also assayed expression of glyceraldehyde-3-phosphate dehydrogenase (GAPDH), a key enzyme in the glycolytic pathway and commonly-used loading control for immunoblots, and found that GAPDH expression was significantly increased in the NR phenotype (Figure 8D). Finally, we assayed expression of the 18kDa translocator protein (TSPO), a constituent of the outer mitochondrial membrane associated with regulation of astrocyte metabolic state^28^, and found that TSPO expression was significantly reduced in LHON-affected astrocytes (Figure 8E).

**Figure 8:**
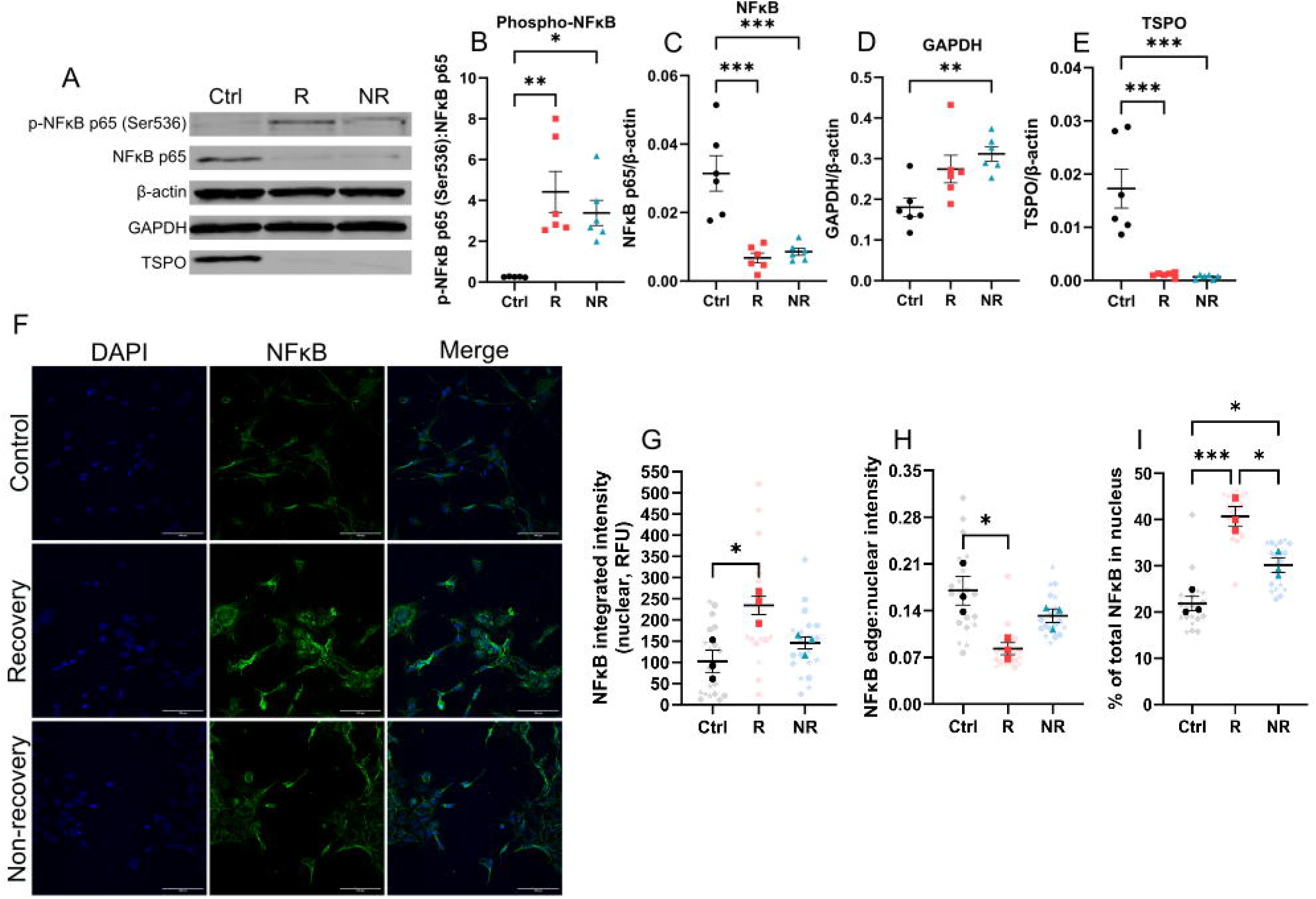
LHON modulated NFκB expression and phosphorylation. A: representative immunoblots. B: NFκB phosphorylation was significantly increased in LHON-affected astrocytes (p_ANOVA_=0.0041, F=8.256, p_Ctrl-R_=0.0037, p_Ctrl-NR_=0.0249, p_R-NR_=0.5709). C: NFκB expression was reduced in LHON affected cells (p_ANOVA_<0.0001, F=18.94, p_Ctrl-R_=0.0002, p_Ctrl-NR_=0.0004, p_R-NR_=0.9148). D: GAPDH expression was increased in LHON-NR astrocytes (p_ANOVA_=0.0076, F=6.864, p_Ctrl-R_=0.0524, p_Ctrl-NR_=0.0070, p_R-NR_=0.5741). E: TSPO expression was significantly reduced in LHON-affected astrocytes (p_ANOVA_<0.0001, F=19.85, p_Ctrl-R_=0.0002, p_Ctrl-NR_=0.0002, p_R-NR_=0.9901). F: immunocytochemical staining of astrocytes for NFκB. G: NFκB integrated intensity was increased in LHON-R (p_ANOVA_=0.0135, F=9.606, p_Ctrl-R_=0.0121, p_Ctrl-NR_=0.3947, p_R-NR_=0.0623). G: nuclear NFκB intensity was increased in LHON-R astrocytes (p_ANOVA_=0.0135, F=9.606, p_Ctrl-R_=0.0121, p_Ctrl-NR_=0.3947, p_R-NR_=0.0623). H: the ratio of nuclear:edge intensity of NFκB was reduced in LHON-R astrocytes (p_ANOVA_=0.0163, F=8.841, p_Ctrl-R_=0.0135, p_Ctrl-NR_=0.2428, p_R-NR_=0.1195). I: the percentage of NFκB colocalised with the nucleus was increased in LHON-affected astrocytes (p_ANOVA_=0.0008, F=29.15, p_Ctrl-R_=0.0007, p_Ctrl-NR_=0.0343, p_R-NR_=0.0128). n=5-6 wells (B-E), n=3 wells (F-I). Data are expressed as mean±SEM. Ordinary one-way ANOVA with Tukey’s multiple comparisons test (B-E). *p<0.05, **p<0.01, ***p<0.001. Scale bars: 100µm.

Next, immunocytochemical staining for NFκB was undertaken (Figure 8F) to confirm nuclear translocation and thus function of NFκB^43^. In line with our immunoblotting data showing increased NFκB phosphorylation at Ser536, we found that LHON-R astrocytes showed a significant increase the integrated intensity of nuclear NFκB (Figure 8G), concomitant with a reduction in the ratio of edge:nuclear NFκB integrated intensity (Figure 8H). Though there was no significant change in total NFκB integrated intensity in LHON astrocytes (Supplementary Figure 9), there was a significant increase in the percentage of NFκB that colocalised with the nuclei of LHON-affected astrocytes (Figure 8I).

### LHON promoted phenotypically divergent secretion of inflammatory factors

We next sought to confirm the impact of LHON on the astrocyte secretome. We deployed a targeted proteomics approach to assay 101 potential targets using pooled samples of astrocyte-conditioned medium from each of our astrocyte phenotypes. Using this approach, we detected a significant shift to the astrocyte secretome, with drastic changes in the expression of various pro-inflammatory cytokines relative to the healthy control (Figure 9; for full list of putative hits see Supplementary Figure 8). We observed concordant upregulations in the secretion of key inflammatory mediators and modulators of the extracellular matrix including extracellular matrix metalloprotease inducer (EMMPRIN), and downregulated secretion of interferon gamma (IFNγ) and dipeptidyl peptidase-4 (DPP4). We further observed phenotypically divergent shifts in the secretion of interleukin-5 (IL-5), brain-derived neurotrophic factor (BDNF), and granulocyte-macrophage colony-stimulating factor (GM-CSF), suggesting that though commonalities exist in the inflamed status of LHON astrocytes, there may be key differences potentially contributing to recovery status. Moreover, we observed polarised shifts in osteopontin expression, which has been implicated in other neurodegenerative conditions^44–46^ and this modulation may contribute to LHON pathology and visual recovery status^46^.

**Figure 9:**
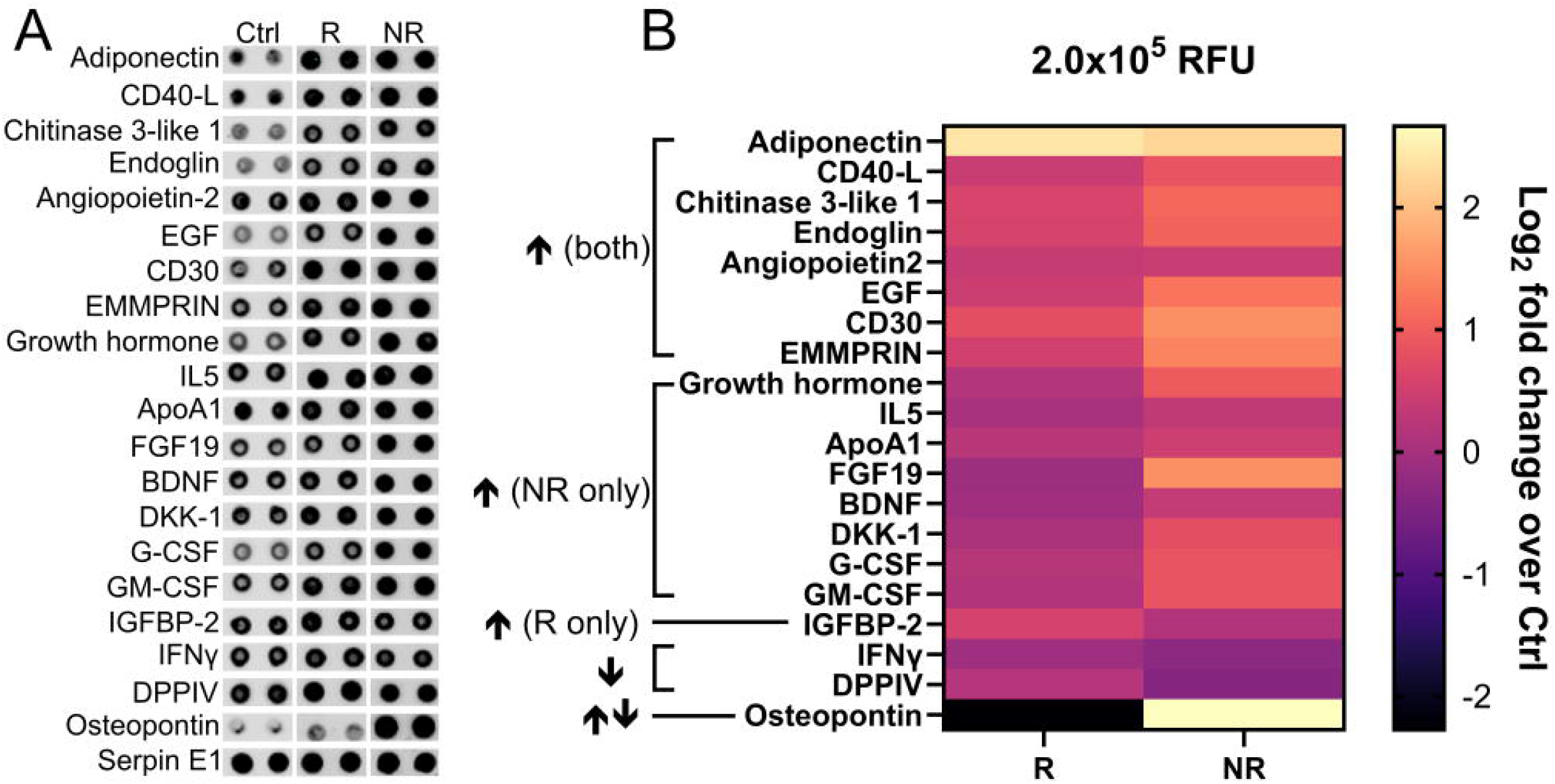
LHON modulates the astrocyte secretome. A: immunoblots of key hits. B: heatmap of key hits expressed as log_2_ fold change over Ctrl. A, B: data are gated to 2.0×10^5^ RFU. n=1 membrane, samples were pooled from 3 independent replicates. CD40-L: cluster of differentiation 40-ligand. EGF: epidermal growth factor. CD30: cluster of differentiation 30. EMMPRIN: extracellular matrix metalloprotease inducer. IL5: interleukin-5. ApoA1: apolipoprotein A1. FGF19: fibroblast growth factor 19. BDNF: Brain-derived neurotrophic factor. DKK-1: Dickkopf-related protein 1. G-CSF: granulocyte-colony stimulating factor. GM-CSF: granulocyte-macrophage colony-stimulating factor. IGFBP-2: insulin-like growth factor binding protein-2. IFNγ: interferon gamma. DPPIV: Dipeptidyl peptidase-4.

## Discussion

Our data represent, to our knowledge, some of the first literature directly addressing the potential role of glial cells in LHON pathology. Our data are suggestive of a phenotypic shift in LHON-affected astrocytes that may help explain both the cascade of RGC degeneration in LHON and the paradoxical nature of spontaneous visual recovery noted in some LHON patients. We focussed on the m.14484T>C genotype because it is associated with the greatest rate of spontaneous visual recovery^5,6,10^ and thus provides a springboard for investigating potential underlying mechanisms in patient-derived samples and data. We report that, regardless of visual recovery status, m.14484T>C-LHON may reduce intracellular levels of GFAP and S100B in iPSC-derived astrocytes, and may enhance GFAP secretion from these same cells. Increased intracellular GFAP is a commonly-reported finding in studies of astrocyte reactivity^35^, however, this outcome is not pan-astrocytic and may not represent all reactive astrocyte phenotypes^23,33–35,47^. Crucially, we observed an upregulation of intra-and extracellular GFAP hyperintensities superficially resembling punctae or aggregates, which may represent diseased astrocyte phenotypes reported in models of other pathological states^23,33,34^. Superficially these findings bear resemblance to clasmatodendrosis in astrocytes^47^ and thus may reflect compromised homeostatic support for retinal ganglion cells; alternatively, the observed GFAP punctae may reflect inappropriate caspase-3 activation in these cells^23,33^. Another potential explanation for the reduction in intracellular GFAP may be that this is a phenomenon associated with mitochondrial dysfunction, as a recent study^48^ described a similar finding of reduced GFAP intensity, aggregation of aldehyde dehydrogenase 1 L 1, and a morphological switch in PINK1-deficient iPSC-derived astrocytes^48^. Regardless of the explanation for this phenomenon, our observations of reduced GFAP (Figure 1A,L; Figure 3A) may help to explain a recent report of enhanced serum GFAP in LHON patients^49^. The superficial resemblance of GFAP puncta to clasmatodendrosis may present a novel avenue for research in LHON tissue, and if possible future work should endeavour to confirm the validity of our observations in LHON patient tissue samples.

Our observation that m.14484T>C-NR LHON may drive AQP4 mislocalisation in astrocytes is also intriguing. AQP4 is a key regulator of water transport in the CNS and its expression is known to increase during astrocyte inflammatory responses^36,37^. Functionally, this may help to explain the transient peripapillary telangiectasia that forms a key hallmark of LHON pathology, with upregulation of AQP4 potentially contributing to this transient swelling. Though AQP4 was increased in LHON astrocytes, its mislocalisation may reflect a lack of function and hence explain the transient swelling, though this must be confirmed in patient tissue samples. We did not assay facets of receptor biology such as receptor internalisation, supramolecular complex formation, or AQP4 activity, so can only speculate about the potential activity status of this increased AQP4 expression. However, a reduction in the ratio of edge:nucleus AQP4 in both LHON phenotypes was observed which may be an additional explanatory factor, as AQP4 expression is associated with astrocyte endfeet under physiological conditions^36^. Secondly, we observed that LHON pathology drove astrocyte swelling and loss of morphological complexity. We suggest that these findings may be linked to mislocalisation of AQP4 to the nucleus, which may also contribute to the alteration of nuclear area we observed in the NR phenotype (which bordered statistical significance, p=0.0512, Figure 1D).

Our observation that LHON did not significantly modulate astrocyte mitochondrial load is potentially unsurprising given recent findings indicating that dysregulated mitophagy:mitobiogenesis underlies pathological status and helps delineate LHON ‘carriers’ from ‘affecteds’^40^. Indeed, enhanced mitophagy or impaired mitobiogenesis may explain why our results trended toward a statistically significant reduction in mitochondrial load, though we did not observe any indication of enhanced mitophagy in our analyses. As a potential alternative explanation, we propose that the lack of effect may be attributable to the astrocyte bioenergetic signature^24^ or the differential organisation of respiratory complexes in astrocytes compared to neurons^50,51^.

Regardless of the effect of LHON on mitochondrial load, we confirmed that LHON phenotypes significantly modulate astrocyte mitochondrial respiration, with the NR phenotype showing a significant deficit in basal mitochondrial respiration. Moreover, both LHON phenotypes showed significant reductions in ATP-linked OCR, which indicates that the mitochondrial oxygen use is less efficient in these cells. These data simultaneously support the historical consensus that LHON reduces cellular ATP production and enhances ROS production^8,38,52^, as ROS production is a feasible explanatory factor for reduced respiratory efficiency as less oxygen would be available for respiration. Antioxidants are emerging as key players in neurodegenerative conditions, including glaucoma^53^, which bears some similarity to LHON pathology and the antioxidant idebenone is currently the only approved LHON therapeutic. Moreover, this may contribute to the inflammatory phenotype we have observed, and exploration of the potential modulatory role of antioxidants on astrocyte bioenergetics and inflammation in LHON should form the basis for future studies. Recent studies in astrocyte immunometabolism implicate a shift toward aerobic glycolysis during the initial inflammatory response, which likely corresponds to the acute phase of inflammation and is accompanied by a pro-inflammatory state^19,28,29^. Seminal works on the contribution of astrocytes to neurodegenerative conditions have suggested that continuous aggravation of the astrocyte inflammatory response without appropriate progression to the resolutory phase of inflammation contributes to the onset of glial asthenia or functional paralysis^54,55^, wherein astrocytes lose the plasticity which forms a core feature of their identity. Consistent with these works, we have similarly observed an increase in aerobic glycolysis concomitant with reduced mitochondrial metabolism in LHON-affected astrocytes, in addition to upregulated phosphorylation of NFκB. We would thus tentatively hypothesise that the Complex I dysfunction associated with LHON pathology, touted to affect ATP and ROS production, may thus contribute to chronic activation of the astrocyte inflammatory response without its appropriate resolution. We have produced evidence in favour of this hypothesis but further work is needed to fully confirm this.

As part of their vital homeostatic function, astrocytes secrete key components of the CNS extracellular matrix (ECM) and perineuronal nets (PNNs)^12,56–58^. Perturbances of this function are associated with astrocyte inflammatory responses and reactivity^59–63^. We observed that LHON pathology was associated with the differential secretion of various inflammatory mediators, including factors known to influence ECM/PNN composition and stiffness. Indeed, ECM degradation and accumulation are factors known to change in various conditions^64,65^, and ECM composition modulates neurite outgrowth^66^, thus providing a promising therapeutic angle for neuroregenerative medicine^67^. In line with this, our finding that osteopontin secretion is polarised in LHON-R and -NR astrocytes respectively may help to explain the capacity for spontaneous visual recovery. Osteopontin is a multifunctional phosphorylated glycoprotein most well-known for its contribution to ossification with further links to cellular migration, inflammatory cytokine production, and modulation of the ECM^68^ that has been implicated in other models of retinal neurodegeneration^44,69^. Indeed, osteopontin expression has been linked glaucomatous and other optic neuropathies^44,46,70,71^, potentially indicating a wider role for neuroinflammation in LHON pathology. In addition to osteopontin, we observed changes to other proteins, including EMPRINN, further supporting a role for ECM dysfunction in LHON pathology. Additionally, in both LHON phenotypes, we observed an increase in secretion of chitinase-3 like protein 1, which is associated with neuroinflammation^72,73^. LHON-NR astrocytes demonstrated a significant increase in GM-CSF secretion, which is associated with the activation of microglia^74–76^, further supporting our suggestion that astrocytes may mediate a neuroinflammatory phenotype in LHON.

Our study has several limitations. Foremost among these are the small sample size, and the lack of female donors. This limits the general applicability of our findings, and we would encourage an expansion on this work by incorporating a larger number of donors and donors of both sexes, age-matched and in balanced numbers. Indeed, the focus on the male sex is a key limiting factor of medical research^77,78^. Moreover, there is a suspected endocrine component to LHON^4,79^ which cannot be fully understood without studying this pathology in both sexes. How paracrine and endocrine secretion of sex-dependent factors such as (neuro)steroid production and metabolism influence the astrocyte phenotype in LHON remain to be studied. It will be important to better understand how our findings are conserved across sexes, as this may limit the therapeutic applicability of our data. Secondly, our study has focussed on one population of glial cells. Although astrocytes comprise a significant body of non-neuronal cells, contributing to 20-40% of the population level depending on the area studied^12^, this does not preclude a potential role for microglia and oligodendrocytes in LHON pathology. To our knowledge, these populations remain to be studied in the context of LHON and we encourage research into these areas to develop a more holistic understanding of this complex pathology. In addition, we have also been limited to an *in vitro* study of astrocyte pathophysiology in LHON, due in part to a lack of a reliable *in vivo* model that accurately recapitulates the causes and progression of this disorder. This is due in part to the complex nature of LHON, in that genetic mutations are necessary but not sufficient to precipitate this condition. Validation of these findings in *post mortem* tissue samples *in absentia* of an *in vivo* model will be required to fully comprehend the clinical validity of this work. Moreover, astrocytes are a highly heterogeneous group of cells^80–82^. Clearly accurate recapitulation of this *in vitro* is a mountainous and perhaps insurmountable task, but use of alternative differentiation methods to yield alternative astrocyte subpopulations (e.g., Müller glia) will be important for further validating these findings. Finally, our work does not address the effect of LHON mutations on astrocyte Ca^2+^ signalling. This is a major signalling modality in astrocytes, and addressing this limitation will aid in the comprehension of these data. Nonetheless, these limitations do not negate the validity or novelty of our work, which provide some of the first direct evidence implicating a role for astrocytes in LHON pathology and as potential influencers of clinical outcomes. Indeed, we believe our findings have significant value and hope that they form the basis for vigorous dissection of glial pathophysiology in the context of LHON.

Future studies should focus on further elucidating the LHON-affected astrocyte metabolic phenotype. In addition, a combined proteomics/transcriptomics screen would be beneficial for more fully elucidating LHON-induced changes to the astrocyte phenotype and LHON-phenotype-specific differences. Astrocytes are a highly heterogeneous group of glia, thus more specific differentiation protocols may yield intriguing results in the absence of a reliable animal model for this pathology to determine subtype-specific contributions to pathological outcomes. Donor sex should also be considered, and a balanced, age-matched approach should be taken to fully elucidate the potential of sexually dimorphic LHON phenotypes.

In summary, using patient-derived materials, we have provided evidence of a potential role for astrocytes in LHON pathology. Taken together, these findings indicate that astrocyte biology is perturbed by LHON pathology, and may in turn influence the course of the disease. Moreover, these data suggest that potential phenotypic differences in astrocyte inflammatory responses may contribute to individual capacity for visual recovery. Our data indicate that RGC pathology in LHON can no longer be considered in isolation, and we urge further study of neuroglial dysfunction in this neurodegenerative condition. Importantly, these findings correlate with findings from other neurological conditions, and may help to explain curious facets of LHON pathophysiology.

## Acknowledgements

The authors would like to thank the funders for facilitating this work, Cardiff University School of Optometry and Vision Sciences Technical Services team for their support, and the iPSC donors for providing the foundational material for this study.

## Author contributions

Wyn Firth: investigation, methodology, study design, funding acquisition, writing – original draft, writing – review and editing, data analysis, data curation. Lubica Dudakova: resources, writing – review and editing. Robert Dobrovolny: resources, writing – review and editing. Tomas Honzik: resources, writing – review and editing. Petra Liskova: funding acquisition, resources, writing – review and editing. Julie Albon: study design, funding acquisition, writing – review and editing. Marcela Votruba: conceptualisation, study design, supervision, funding acquisition, writing – review and editing, data curation.

## Data availability statement

The data that support the findings of this study are available from the corresponding author upon reasonable request.

## Abbreviations

ACC: Acetyl coenzyme A carboxylase
AMPK: Adenosine monophosphate-activated protein kinase
AQP4: Aquaporin-4
ASFMM: Astrocyte serum-free maintenance media
ATP: Adenosine triphosphate
BDNF: Brain-derived neurotrophic factor
CNS: Central nervous system
DPP4/DPPIV: Dipeptidyl peptidase-4
ECM: Extracellular matrix
EMPRINN: Extracellular matrix metalloprotease inducer
GAPDH: Glyceraldehyde-3-phosphate dehydrogenase
GFAP: Glial fibrillary acidic protein
GM-CSF: Granulocyte-macrophage colony-stimulating factor
IFNγ: Interferon gamma
IL-5: Interleukin-5
iPSC(s): induced Pluripotent Stem Cell(s)
LHON: Leber’s Hereditary Optic Neuropathy
NFκB: Nuclear factor kappa-light-chain-enhancer of activated B cells
NPC(s): Neural progenitor cell(s)
NR, LHON-NR: Non-recovery
OCR: Oxygen consumption rate
OXPHOS: Oxidative phosphorylation
PBS: Phosphate-buffered saline
PER: Proton efflux rate
PNN(s): Perineuronal net(s)
R, LHON-R: Recovery
RFU: Relative fluorescence units
RGC(s): Retinal ganglion cell(s)
ROS: Reactive oxygen species
RTP: Room temperature and pressure
S100B: S100 calcium binding protein B
STAT3: Signal transducer and activator of transcription 3
TBS: Tris-buffered saline
TOMM20: Translocase of the Outer Mitochondrial Membrane 20
TSPO: Translocator protein 18kDa

