## Supplementary Figures for "Bioenergetic dysfunction and inflammation in hiPSC-derived astrocytes from m.14484T>C Leber’s Hereditary Optic Neuropathy"

Wyn Firth[1*] (ORCID: **0000-0001-6531-2580**), Lubica Dudakova[2] (ORCID: **0000-0003-4718-8955**), Robert Dobrovolny[2] (ORCID: **0000-0003-0081-156X**), Tomas Honzik[2] (ORCID: **0000-0003-4300-2519**), Petra Liskova[2,3] (ORCID: **0000-0001-7834-8486**), Julie Albon[1,4] (ORCID: **0000-0002-3029-8245**), Marcela Votruba*[1, 5] (ORCID: **0000-0002-7680-9135**)

### Supplementary Figure 1

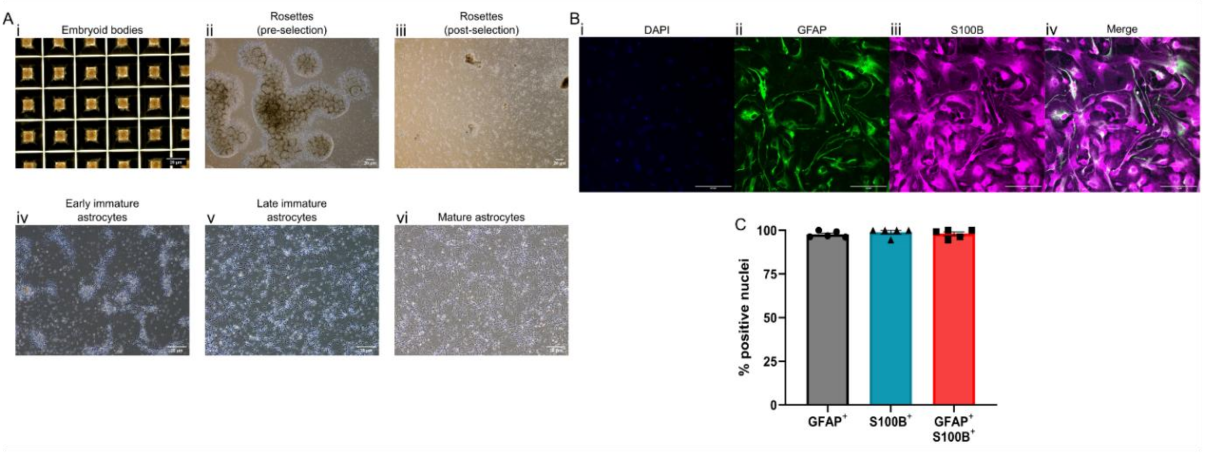


Astrocyte differentiation. Ai-vi: representative images showing differentiation from iPSC-based embryoid bodies (i) to mature astrocytes (vi) over the course of 60 days. B: confirmation of astrocyte fate specification. i: nuclear counterstain DAPI. ii: GFAP staining. iii: S100B staining. iv: merge of all channels. C: quantification of astrocyte marker expression; datapoints are 5 fields of view from 1 coverslip. Cells were 100% positive for GFAP/S100B expression or coexpression. Scale bars: 20μm (A), 100 μm (B).

### Supplementary Figure 2

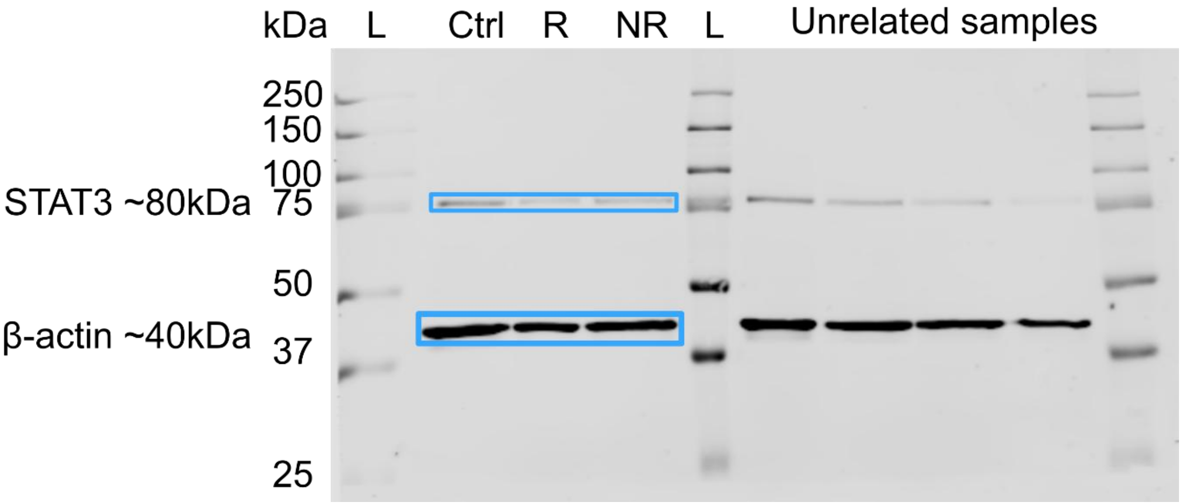


Full uncropped immunoblot related to Figure 7. Blue boxes show regions of interest for bands indicated.

### Supplementary Figure 3

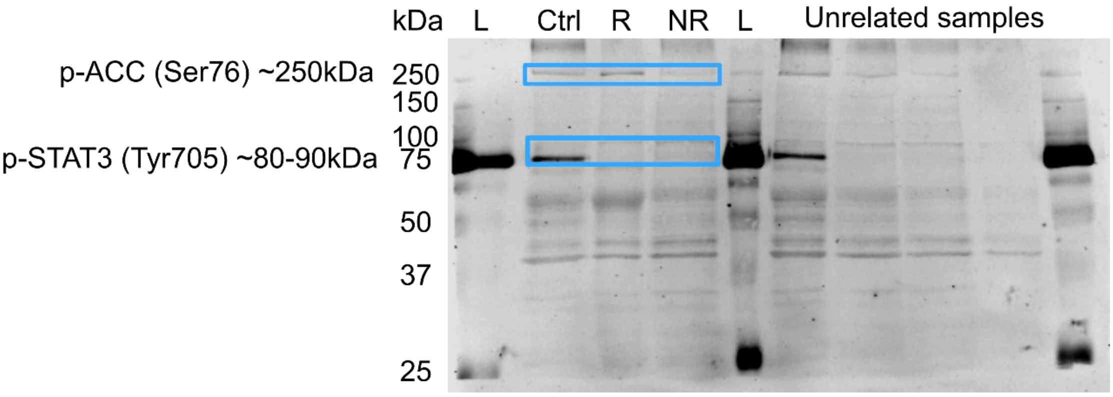


Full uncropped immunoblot related to Figure 7. Blue boxes show regions of interest for bands indicated.

### Supplementary Figure 4

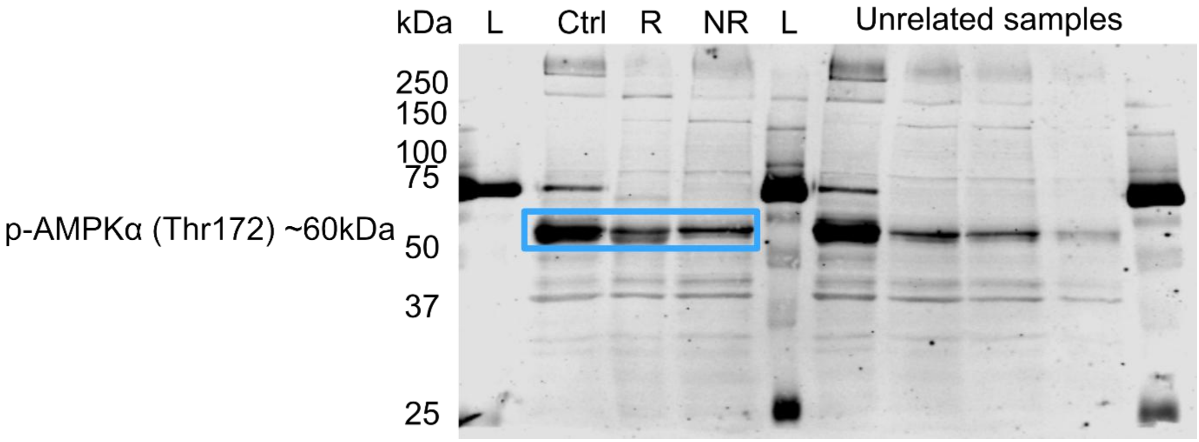


Full uncropped immunoblot related to Figure 7. Blue boxes show regions of interest for bands indicated.

### Supplementary Figure 5

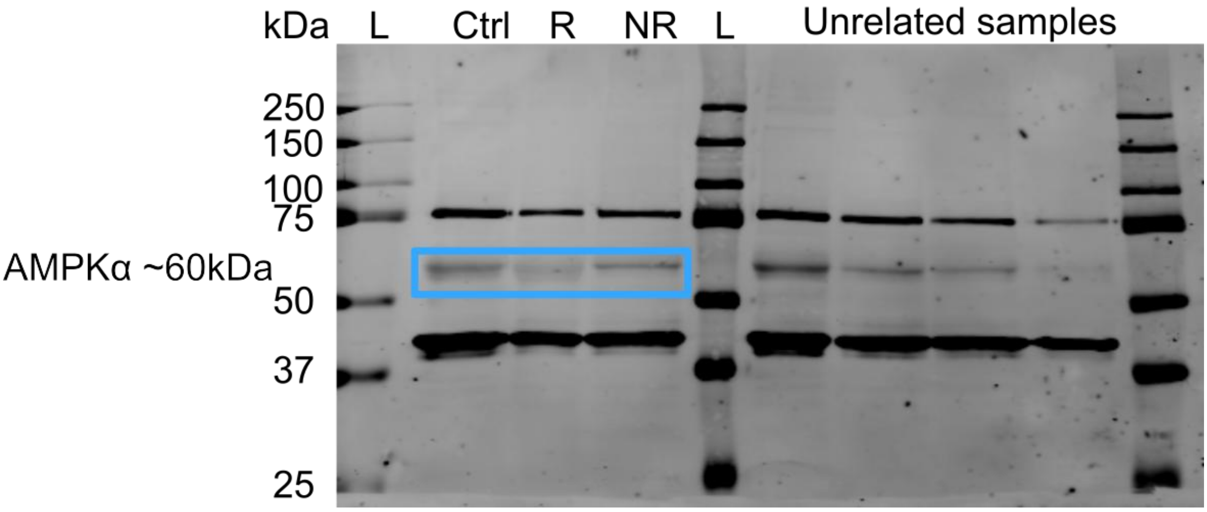


Full uncropped immunoblot related to Figure 7. Blue boxes show regions of interest for bands indicated.

### Supplementary Figure 6

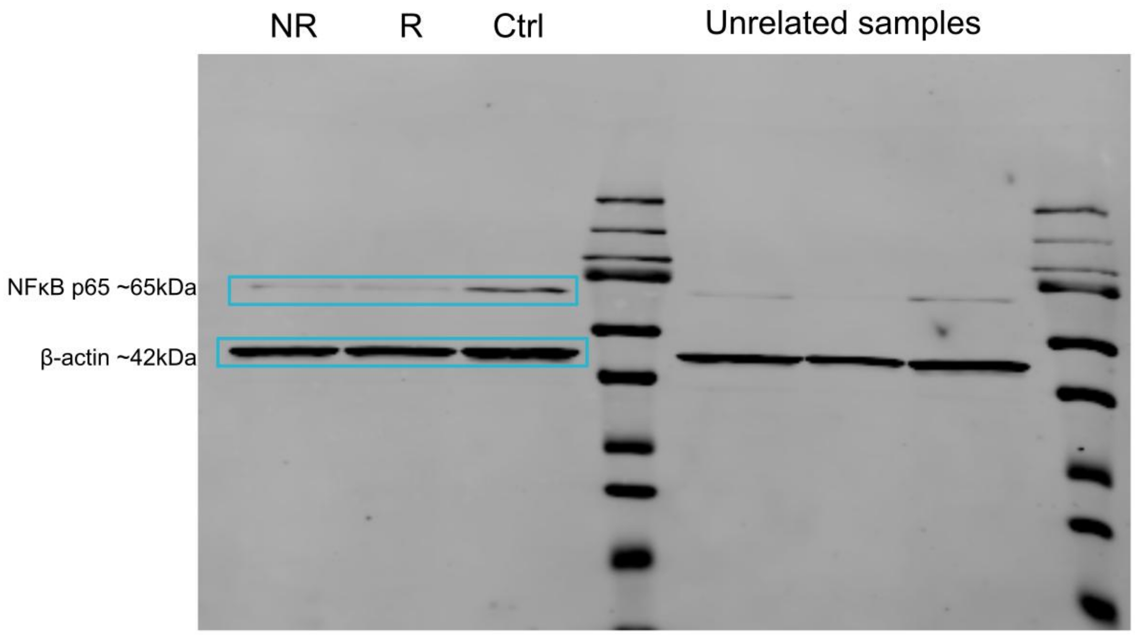


Full uncropped immunoblot related to Figure 8. Blue boxes show regions of interest for bands indicated.

### Supplementary Figure 7

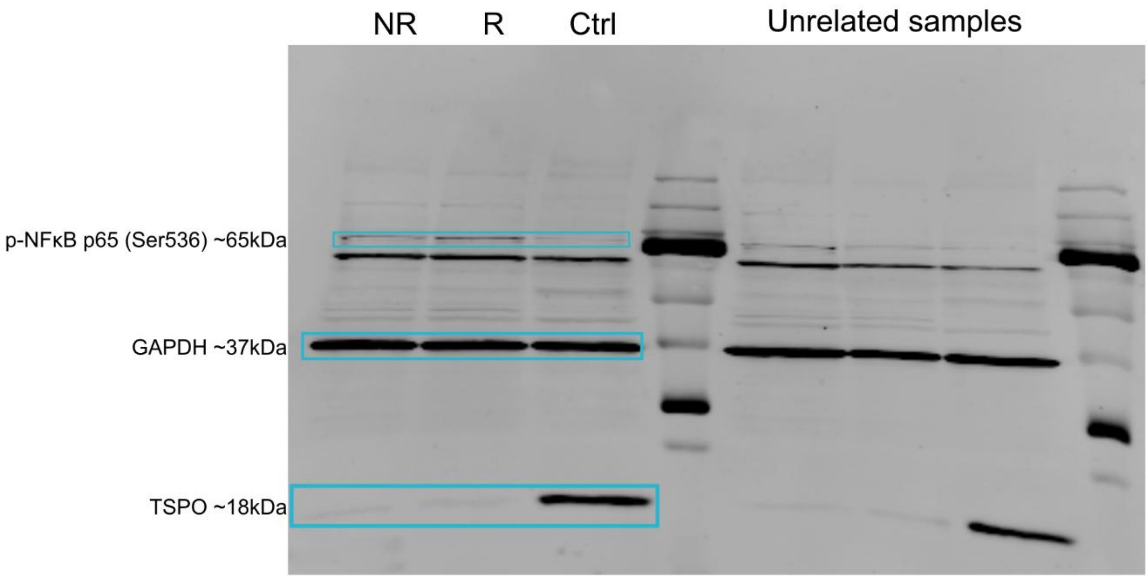


Full uncropped immunoblot related to Figure 8. Blue boxes show regions of interest for bands indicated.

### Supplementary Figure 8

A

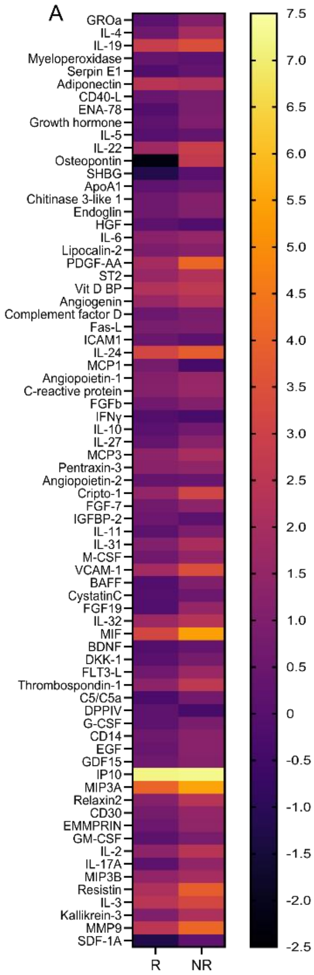


B

| Target | Group | | |
|---|---|---|---|
| | Ctrl | R | NR |
| Leptin | 1 | 28255 | 34640 |
| LIF | 1 | 17205 | 26690 |
| IL23 | 1 | 15655 | 32290 |
| IL8 | 1 | 17455 | 28040 |
| PDGF-AB/BB | 1 | 7755 | 15690 |
| TARC | 1 | 14105 | 30890 |
| CD31 | 1 | 24255 | 36240 |
| TFF3 | 1 | 19255 | 19490 |
| TIM3 | 1 | 26105 | 42740 |
| PF4 | 1 | 12905 | 31940 |
| TfR | 1 | 20905 | 68840 |
| IGF-BP3 | 1 | 19055 | 36340 |
| IL-12 P70 | 1 | 23155 | 44140 |
| RAGE | 1 | 20955 | 46240 |
| TGFa | 1 | 20805 | 46240 |
| IL-1A | 1 | 16905 | 51990 |
| IL-13 | 1 | 4855 | 22340 |
| IL-33 | 1 | 8755 | 27740 |
| MIG | 1 | 22205 | 43840 |
| RANTES | 1 | 11455 | 30140 |
| IL-1B | 1 | 15205 | 31690 |
| IL-15 | 1 | 20205 | 41990 |
| IL-34 | 1 | 12955 | 29240 |
| MIP1A/B | 1 | 14055 | 30890 |
| RBP4 | 1 | 19205 | 37940 |
| TNF | 1 | 12055 | 26490 |
| IL-1RA | 1 | 24005 | 47590 |
| IL-16 | 1 | 13805 | 46390 |
| uPAR | 1 | 19255 | 38690 |
| VEGF | 1 | 9905 | 34240 |
| IL-18 BPA | 1 | 17605 | 44940 |

Expression (relative fluorescence units)

C

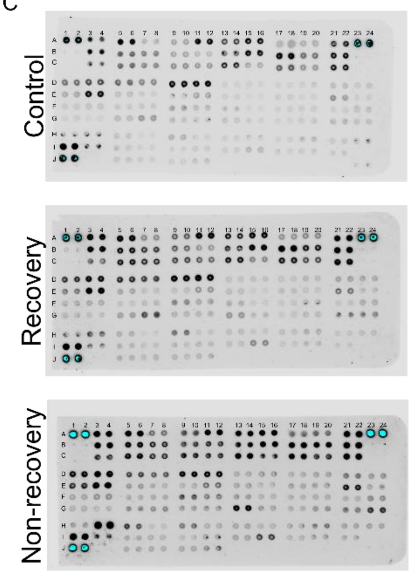


Full list of hits from Human XL Cytokine Array (BioTechne, ary022b), related to Figure 9. Control values that were undetectable have been termed ‘1’ for the purpose of relative expression. A: heatmap showing $log_2$ fold change of proteins over healthy control where healthy control expression is detectable. B: table showing list of hits where expression in healthy control was below the limit of detection. C: full uncropped membranes.

### Supplementary Figure 9

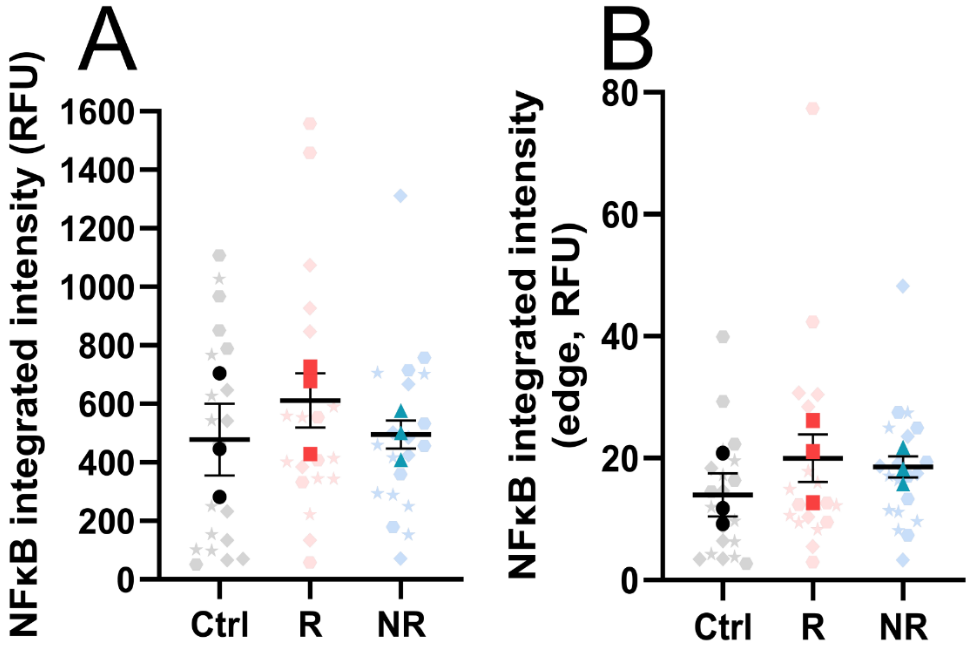


NFκB immunocytochemical staining results, related to Figure 8.
